# Neuroscience Cloud Analysis As a Service

**DOI:** 10.1101/2020.06.11.146746

**Authors:** Taiga Abe, Ian Kinsella, Shreya Saxena, E. Kelly Buchanan, Joao Couto, John Briggs, Sian Lee Kitt, Ryan Glassman, John Zhou, Liam Paninski, John P. Cunningham

**Affiliations:** Mortimer B. Zuckerman Mind Brain Behavior Institute; Center for Theoretical Neuroscience, Columbia University; Grossman Center for the Statistics of Mind, Columbia University; Department of Neuroscience, Columbia University; Department of Statistics, Columbia University; Department of Computer Science, Columbia University; Department of Neurobiology, University of California

## Abstract

A major goal of computational neuroscience is the development of powerful data analyses that operate on large datasets. These analyses form an essential toolset to derive scientific insights from new experiments. Unfortunately, a major obstacle currently impedes progress: novel data analyses have a hidden dependence upon complex computing infrastructure (e.g. software dependencies, hardware), acting as an unaddressed deterrent to potential analysis users. While existing analyses are increasingly shared as open source software, the infrastructure needed to deploy these analyses – at scale, reproducibly, cheaply, and quickly – remains totally inaccessible to all but a minority of expert users. In this work we develop Neuroscience Cloud Analysis As a Service (NeuroCAAS): a fully automated analysis platform that makes state-of-the-art data analysis tools accessible to the neuroscience community. Based on modern large-scale computing advances, NeuroCAAS is an open source platform with a drag-and-drop interface, entirely removing the burden of infrastructure purchase, configuration, deployment, and maintenance from analysis users and developers alike. NeuroCAAS offers two major scientific benefits to any data analysis. First, NeuroCAAS provides automatic reproducibility of analyses at no extra effort to the analysis developer or user. Second, NeuroCAAS cleanly separates tool implementation from usage, allowing for immediate use of arbitrarily complex analyses, at scale. We show how these benefits drive the design of simpler, more powerful data analyses. Furthermore, we show that many popular data analysis tools offered through NeuroCAAS outperform typical analysis solutions (in terms of speed and cost) while improving ease of use, dispelling the myth that cloud compute is prohibitively expensive and technically inaccessible. By removing barriers to fast, efficient cloud computation, NeuroCAAS can dramatically accelerate both the dissemination and the effective use of cutting-edge analysis tools for neuroscientific discovery.

## 1 Introduction

Driven by the constant evolution of new recording technologies and the vast quantities of data that they generate, neural data analysis — which aims to build the path from these datasets to scientific insight — has grown into a centrally important part of modern neuroscience, enabling significant new insights into the relationships between neural activity, behavior, and the external environment (Paninski and Cunningham, 2018). As a key part of this growth, the complexity of neural data analyses has massively increased. Historically, the software implementation of a data analysis (what we call the core analysis- Figure 1A) was typically a small, isolated code script with few dependencies. In stark contrast, modern core analyses routinely incorporate video processing algorithms (Pnevmatikakis et al., 2016, Pachitariu et al., 2017, Mathis et al., 2018, Zhou et al., 2018, Giovannucci et al., 2019), deep neural networks (Batty et al., 2016, Gao et al., 2016, Lee et al., 2017, Parthasarathy et al., 2017, Mathis et al., 2018, Pandarinath et al., 2018, Giovannucci et al., 2019), sophisticated graphical models (Yu et al., 2009, Wiltschko et al., 2015, Gao et al., 2016, Wu et al., 2020), and other cutting-edge machine learning tools (Pachitariu et al., 2016, Lee et al., 2017).

**Figure 1:**
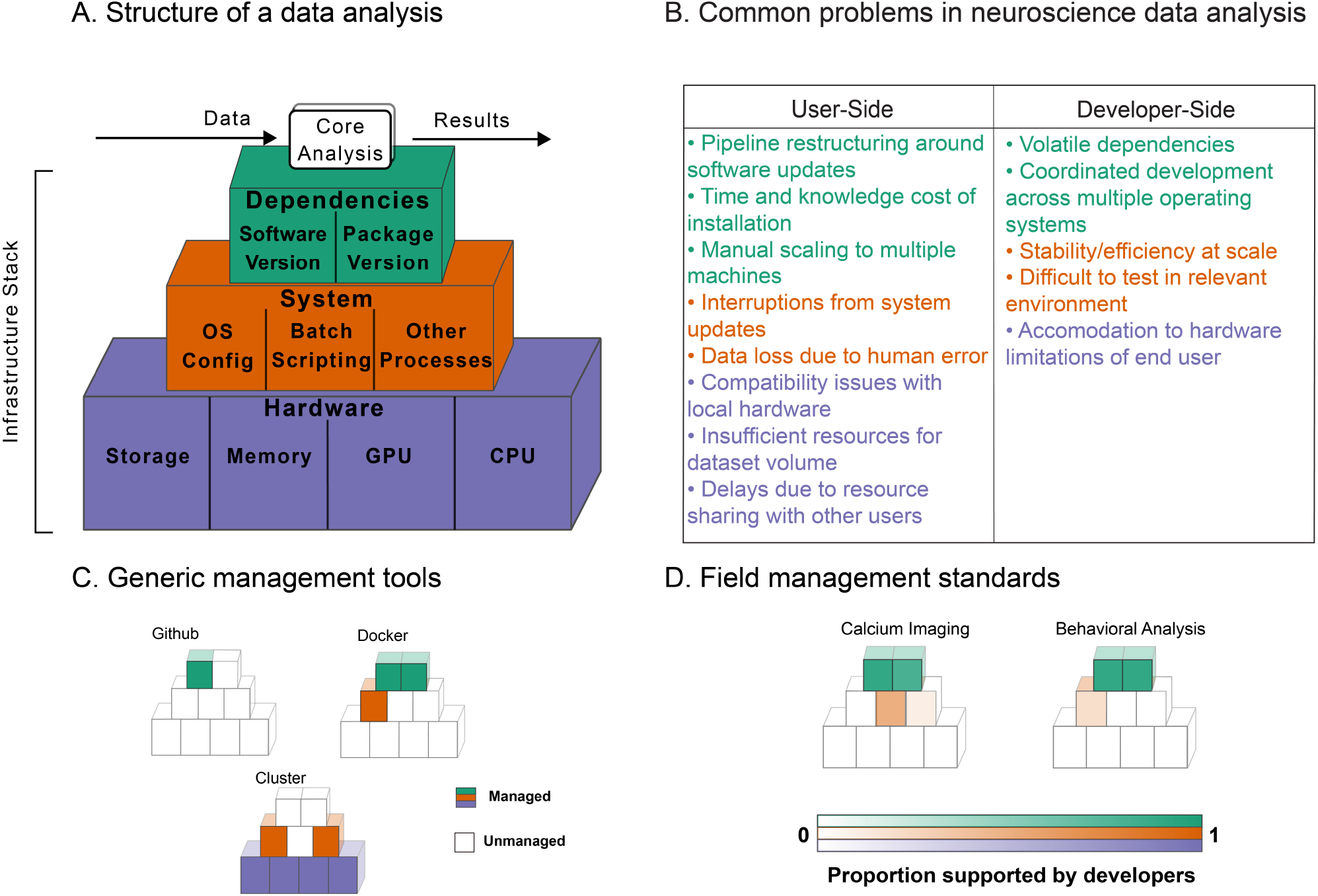
Data analysis infrastucture. A. Core analysis code sits atop an infrastructure stack (including both software and hardware) that must be installed and maintained to make analysis viable. B. A number of common problems arise at each layer of this infrastructure stack; some of these issues are most visible to users, while some are more visible to software developers. C. Many common management tools deal only with one or two layers in the infrastructure stack, leaving gaps that users and developers must fill manually or by cobbling systems together. D. Our surveys of common neural data analysis tools for calcium imaging and behavioral analysis indicate that many layers of the infrastructure stack are currently not managed by analysis developers, and implicitly delegated to the user’s responsibility (see **Materials and Methods** for full details).

Importantly, however, as developers build more powerful analyses, core analysis software becomes increasingly coupled to underlying analysis *infrastructure* (Figure 1A): software dependencies like the deep learning libraries PyTorch and TensorFlow (Abadi et al., 2016, Paszke et al., 2019), system level dependencies to manage jobs and computing resources (Merkel, 2014), and hardware dependencies such as a precisely configured CPU, access to a GPU, or a required minimum amount of device memory. Figure 1A details the way these individual components comprise a full infrastructure *stack*: the necessary – but largely ignored – foundation of resources on which all data analyses run (Demchenko et al., 2013, Jararweh et al., 2016, Zhou et al., 2016).

The immediate implications of neglected infrastructure are a set of problems all too familiar to the neuroscience community: for every novel analysis, labor and financial resources must be spent on hardware setup, software troubleshooting, unexpected interruptions during long analysis runs, processing constraints due to limited “on-premises” computational resources, and more (Figure 1B). However, far from simply being a nuisance, neglected infrastructure has wide reaching and urgent scientific consequences. Most prominently, unaddressed infrastructure impacts analysis reproducibility. As data analyses become more dependent on specific, complex infrastructure stacks, it becomes extremely difficult for developers to document and maintain reproducible analyses (Monajemi et al., 2019, Nowogrodzki, 2019). It has been noted that the current state of analysis infrastructure is a major contributor to the endemic lack of reproducibility suffered by modern data analysis (Crook et al., 2013, Hinsen, 2015, Stodden et al., 2018, Krafczyk et al., 2019, Raff, 2019), and that infrastructure-based barriers are an unaddressed obstacle to the proliferation of innovative analyses (Magland et al., 2020). Specific cases where infrastructure issues have culminated in critical misrepresentations of big datasets have been well documented in adjacent fields (Miller, 2006, Glatard et al., 2015), and similar challenges have been noted in machine learning, where deep learning specific infrastructure components can dictate model performance (Radiuk, 2017). Illustratively, a recent study of the machine learning literature observed that although local compute clusters claim to address the issue of hardware availability, *none* of the studies that required use of a compute cluster were reproducible (Raff, 2019).

Major efforts have been made by journals (Donoho, 2010, Hanson et al., 2011, “Code Share”, 2014) and funding agencies (Carver et al., 2018) to encourage the sharing of core analysis software. Additionally, new tools have been developed to address scientific challenges like the formatting (Teeters et al., 2015) and storage of data (Dandi Team, 2019), or workflow management on existing infrastructure (Yatsenko et al., 2015, Gorgolewski et al., 2011) (see Platform and Discussion for a detailed overview). However, these important efforts still ignore the majority of the analysis infrastructure stack. Despite calls to improve standards of practice in the field (Vogelstein et al., 2016), and work in fields such as astronomy, genomics, and high energy physics (Hoffa et al., 2008, Zhou et al., 2016, Chen et al., 2019, Monajemi et al., 2019), there has been little concrete progress in our field towards a scientifically acceptable infrastructure solution. Some tools – compute clusters, versioning tools like Github (https://github.com), and containerization services like Docker (Merkel, 2014) – provide various infrastructure components (Figure 1C), but it is nontrivial to combine these components into a complete infrastructure stack. The ultimate effect of these partially used toolsets is a hodgepodge of often slipshod infrastructure practices (Figure 1D; see supporting data in Tables S1, S2).

Critically, management of these issues most often falls on trainees who are neither scientifically rewarded (Landhuis, 2017, Chan Zuckerberg Initiative, 2014), nor specifically instructed (Merali, 2010) on how to build, configure, and install infrastructure stacks. We term this conventional model *Infrastructure-as-Graduate-Student* (IaGS)– infrastructure treated as a scientific afterthought, addressed only with patchwork tools and operating as a silent source of errors and inefficency. The IaGS status quo fails any reasonable standard of scientific rigor, reduces the accessibility of valuable analytical tools, and impedes scientific training and progress.

Of course, infrastructure challenges are not specific to neuroscience, or even science generally. Entities that deploy software at industrial scale have been empowered in recent years by the *Infrastructure-as-Code* (IaC) paradigm: an emerging toolset that ensures the stability and replicability of infrastructure stacks by automating their creation and management (Morris, 2016, Aguiar et al., 2018). An IaC platform first requires a code document that completely specifies each infrastructure component (each colored block in Figure 1A) supporting any given core software. From this code document, the corresponding infrastructure stack can be activated and assembled automatically, in a process called *deployment.* After deployment, anyone with access to the platform can use the core software in question without ever having touched its underlying infrastructure stack. Users and developers also have the assurance that core software is functioning exactly as indicated in the corresponding code document, skirting all of the issues shown in Figure 1B. However, despite these benefits, there has been no previous effort to extend IaC to neuroscience data analyses and their associated infrastructure stacks.

In response, we developed Neuroscience Cloud Analysis as a Service (NeuroCAAS), an IaC platform that pairs core analyses with bespoke infrastructure stacks (i.e. all the components in Figure 1A) in a deployable code document. In particular, the infrastructure stack is treated as a set of modular components (§2.1.1-2.1.3) corresponding to versions of the software, system, and hardware infrastructure components that can be concisely specified in code. NeuroCAAS then stores the specification of this complete data analysis in a document called a *blueprint* (§2.1.4), which can be deployed to analyze data on demand. To maximize the scale and accessibility benefits of our platform, we provide a open source web interface to NeuroCAAS (§2.2), available to the neuroscience community at large. The result is drag-and-drop usage of neural data analysis: experimentalists and computational neuroscientists can log on to the NeuroCAAS website, set some parameters for an analysis, and simply submit their neural or behavioral data. A new infrastructure stack is then automatically deployed on the cloud according to a specified blueprint and autonomously used to produce analysis results, which are returned to the user. This aspect of NeuroCAAS warrants emphasis, as it diverges starkly from traditional scientific practice: NeuroCAAS is *not* a platform design or suggestion that the reader can attempt to recreate on their own; instead, NeuroCAAS is offered as an open source infrastructure platform available for immediate use, via a website (www.neurocaas.org).

Below, we first outline the structure of the NeuroCAAS platform. Next, we present two example analyses that are enabled by NeuroCAAS’s design, concerning widefield calcium imaging (WFCI) and markerless tracking models for behavioral video. Finally, we quantify the performance of popular neuroscience analyses on NeuroCAAS, and find that analyses encoded on NeuroCAAS are cheaper and faster than analogues run on on-premises infrastructure (e.g. a university compute cluster), in addition to the major benefits in reproducibility, accessibility, and scalability emphasized above.

## 2 Platform

NeuroCAAS is fundamentally a platform that links a collection of customizable infrastructure components (§2.1.1-2.1.3) to a library of analysis blueprints (§2.1.4): each blueprint specifies a standardized configuration of infrastructure components that can be deployed automatically.

To process their data, users simply log in to the platform and drag and drop their dataset(s) into the upload area of a given analysis (Figure 2, top left). After choosing a set of parameters to apply to this data, they submit their data for processing, invoking a NeuroCAAS “job.” No further user input is needed: given the relevant dataset and parameters, NeuroCAAS sets up the requested analysis on a new infrastructure stack from the corresponding blueprint (Figure 2, black arrows) and analyzes data on this infrastructure (Figure 2, blue arrows), providing scalable and reproducible computational processing on demand (Figure 2, bottom right). Analysis outputs (including live status logs and a complete description of analysis parameters) are then delivered back to the user (Figure 2, bottom left), and finally all created infrastructure is dissolved when data processing is complete (Figure 2, bottom right). As an example, the video in Figure 3 shows how users can submit data and track the training of three separate DeepGraphPose models (Wu et al., 2020) simultaneously on NeuroCAAS.

**Figure 2:**
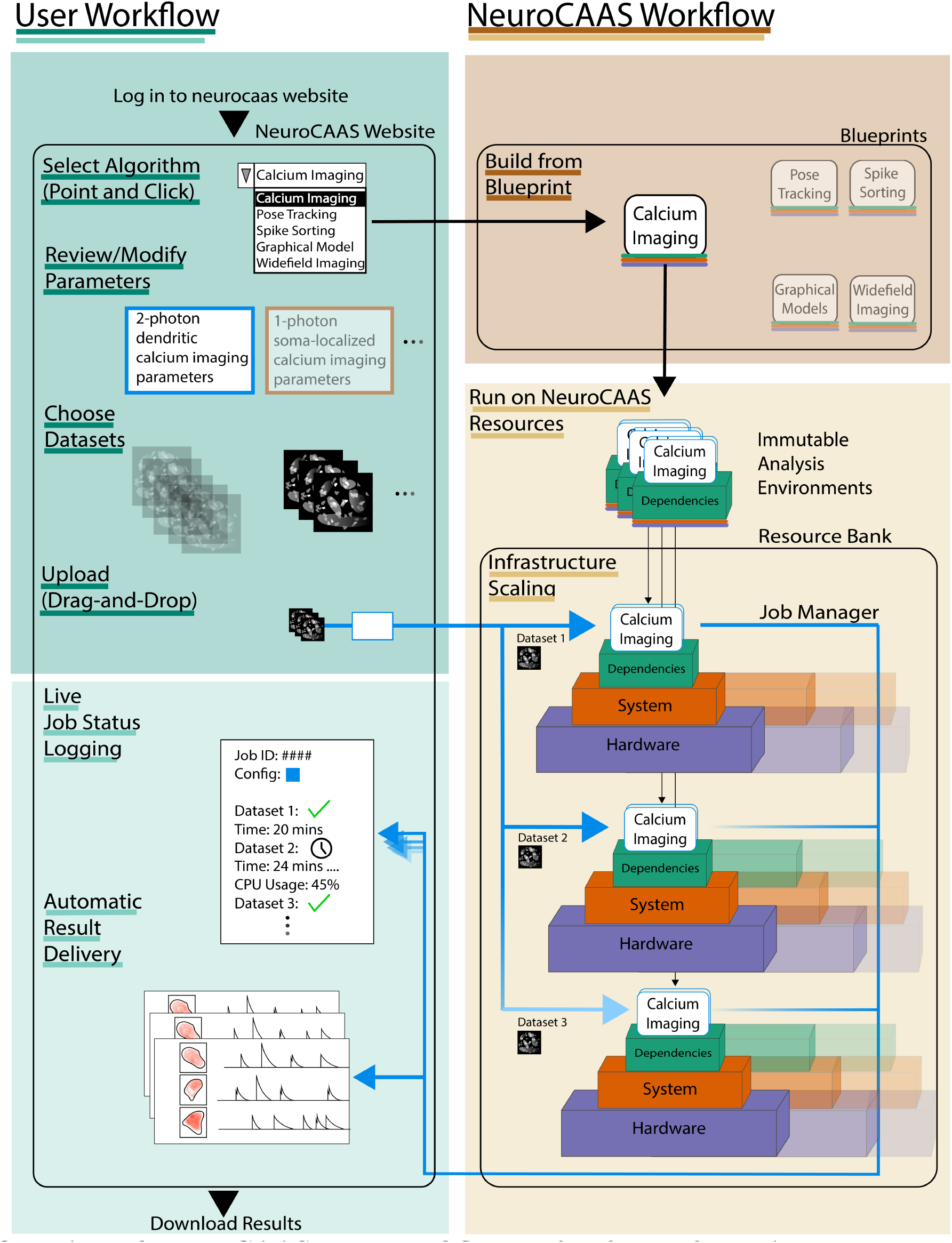
Overview of NeuroCAAS User Workflow. Left indicates the user’s experience; right indicates the work that NeuroCAAS performs. The user begins by choosing from the analysis encoded in NeuroCAAS. They then modify corresponding configuration parameters as needed. Finally, the user uploads dataset(s) and a configuration file for analysis. NeuroCAAS detects this upload event and deploys the requested analysis using an infrastructure blueprint. NeuroCAAS builds the appropriate number of immutable analysis environments and sets up corresponding resources. Multiple analysis environments may be deployed in parallel if the user uploads multiple datasets- and the job manager automatically handles input and output scaling. The deployed resources persist only as necessary, and results, as well as diagnostic information, are automatically routed back to the user.

**Figure 3:**
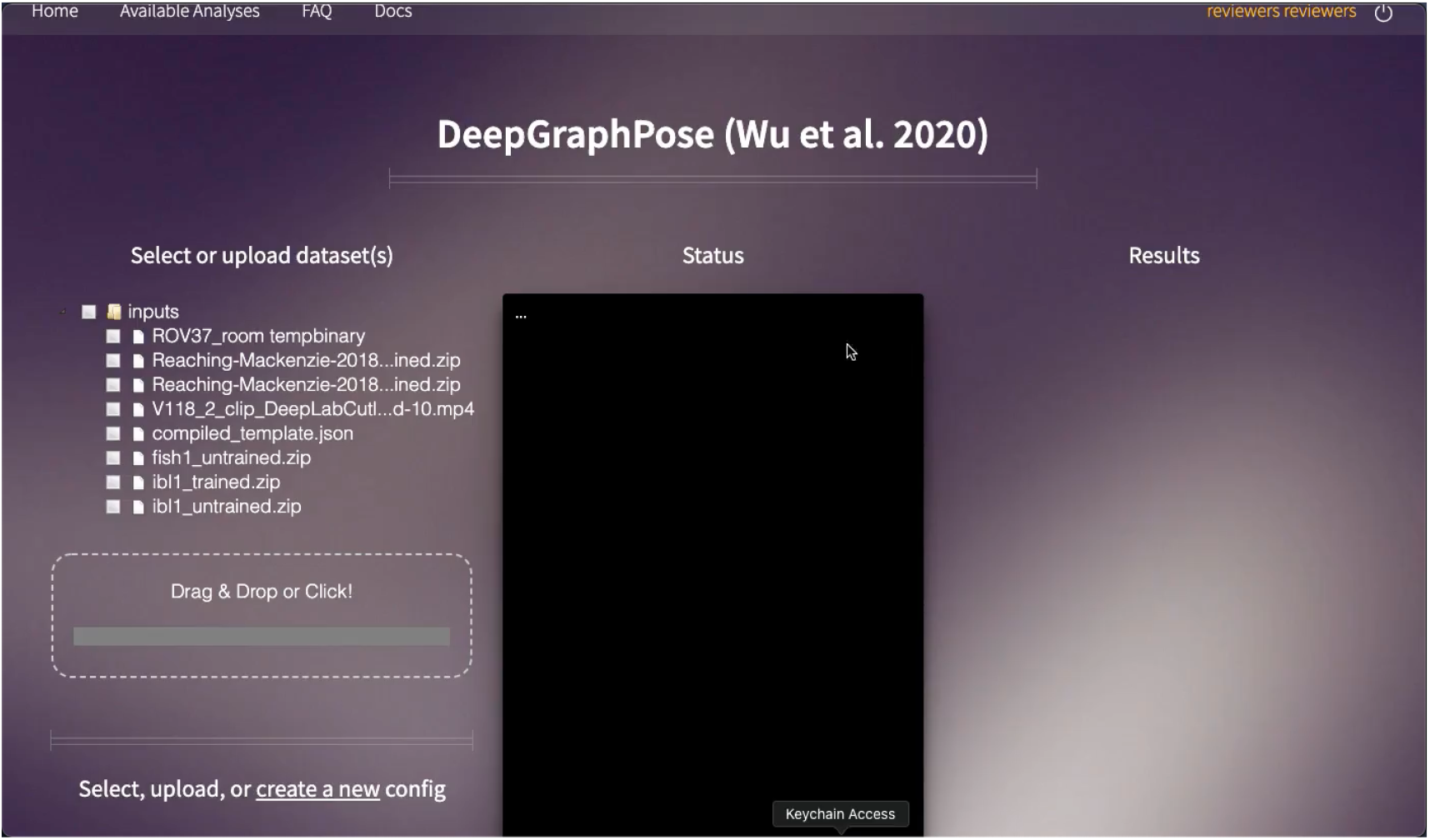
Example workflow run on NeuroCAAS website. Demonstration run of DeepGraphPose (Wu et al., 2020) training three separate models at the same time. This is a truncated run where the model is trained for only a few iterations on each step of DeepGraphPose training (for details see Wu et al. (2020)). Caption text at screen bottom explains each step of processing. Video is dgp-train-triple-fulldemo.mp4.

In what follows, we will first describe each part of the NeuroCAAS platform in §2.1. We will then discuss the interface to the NeuroCAAS platform in section §2.2. Finally, we survey and contrast related systems in §2.3.

### 2.1 NeuroCAAS Structure

The structure of NeuroCAAS naturally circumvents the issues of reproducibility, accessibility, and scale that burden existing infrastructure tools and platforms. NeuroCAAS breaks the infrastructure stack into three decoupled parts that together are sufficient to support virtually any given core analysis. Critically, the configuration of each of these parts is designed to be concisely summarizable in a blueprint (§2.1.4). First, to address all software level infrastructure, NeuroCAAS offers all analyses as *immutable* analysis environments (§2.1.1). Second, to address system configuration, each NeuroCAAS analysis has a built-in job manager (§2.1.2) that automates all of the logistical tasks associated with analyzing data: configuring hardware, logging outputs, parallelizing jobs and more. Third, to provide specific computing hardware on demand, NeuroCAAS manages a resource bank (§2.1.3) of hardware and operating systems built on the public cloud, making the service globally accessible and effectively limitless. We describe the value of each of these components in depth in §5.1.

#### 2.1.1 Immutable analysis environments for software infrastructure

To make software infrastructure concisely summarizable in code, NeuroCAAS serves analyses to users exclusively through immutable analysis environments (IAEs). An IAE is an isolated environment containing the installed core analysis code and all necessary software dependencies, similar to a Docker container (Merkel, 2014). An IAE also contains a workflow script that parses input and parameters in a prescribed way and runs the steps of a given analysis (Figure 2, right; Figure 5, top left). In stipulating immutability (data analysis cannot be altered once started), IAEs automate analysis installation and workflow, eliminating the possibility of bugs resulting from incompatible dependencies, mid-analysis misconfiguration (Figure 4A, installation and troubleshooting), or other so-called “user degrees of freedom” and ensure that analyses are run within *developer-defined* parameters. Immutability has a long history as a principle of effective programming and resource management in computer science (Bloch, 2008, Morris, 2016), and in this context is closely related to the view that data analysis should be automated as much as possible (Tukey, 1962, Waltz and Buchanan, 2009). These views are motivated by the observation that immutability and automation allow applications to scale easily as they become more complex, a feature we make use of in our NeuroCAAS native analyses.

**Figure 4:**
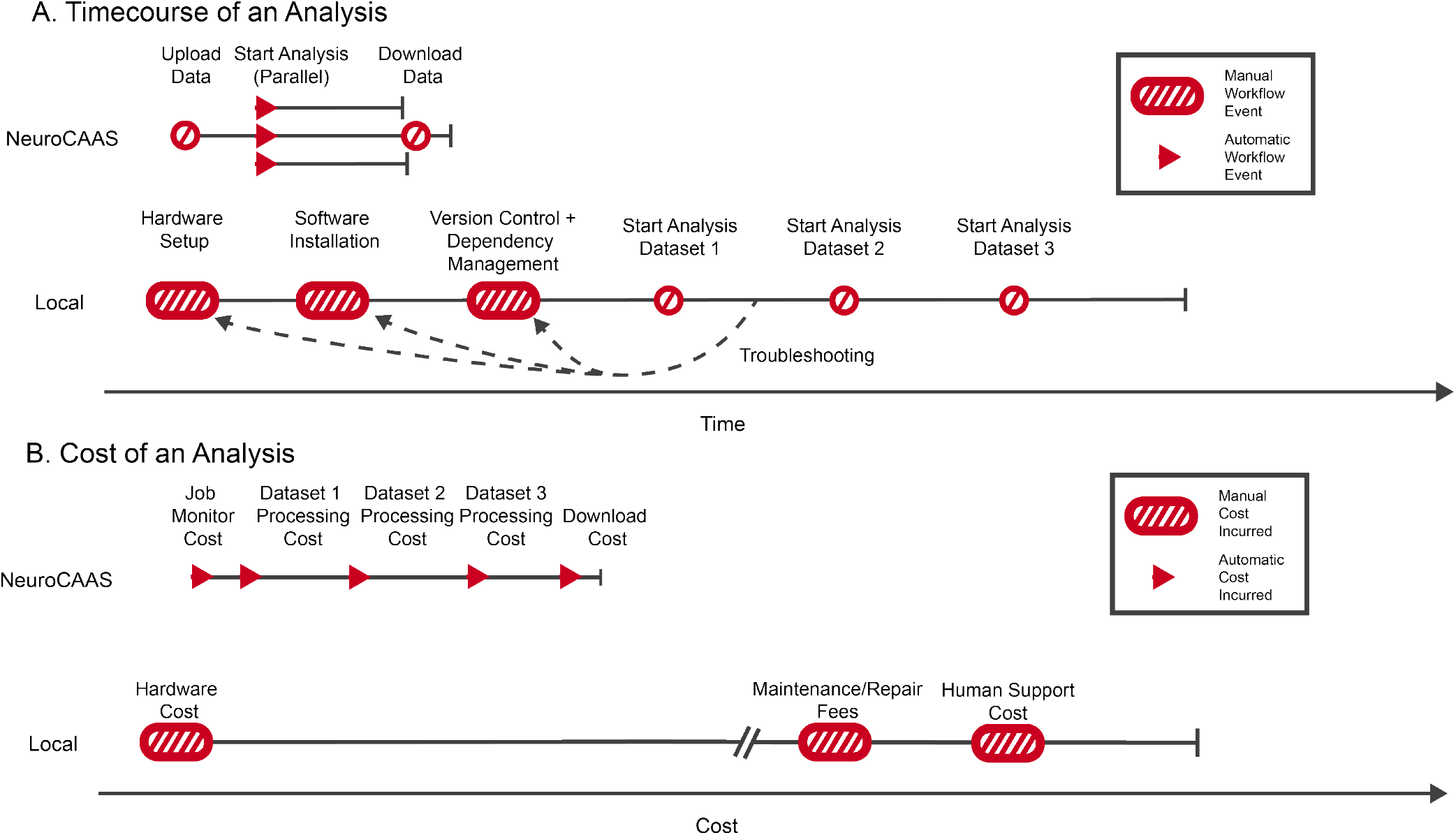
Infrastructure-as-grad-student vs infrastructure-as-code. A. Local processing via IaGS requires a number of time-consuming steps from the user (hardware setup, software installation and maintenance, etc.) before any analyses are run. Then typically analyses of large datasets are run serially (due to resource constraints), leading to longer processing times. On NeuroCAAS, user interaction is only required at the beginning of the analysis (to upload the data), then NeuroCAAS processes the data using large-scale parallel compute resources, leading to faster overall processing times. B. On NeuroCAAS, some costs are incurred with each analysis run: the user must upload the datasets (incurring a small job monitor cost), and then each dataset incurs some compute cost. For local processing, the bulk of the costs are paid upfront, in purchasing hardware; then additional labor costs are incurred for maintenance, support, and usage of limited local resources. If the per-dataset costs are low and the total number of datasets to be processed is limited then NeuroCAAS can lead to significantly smaller total costs than local processing.

Each IAE has a unique ID from which it can be set up on almost any computer, transforming the computer into an input-output machine for the corresponding analysis. We have currently implemented 13 different analyses in immutable analysis environments (Table 1).

**Table 1:** List of existing analyses currently implemented on NeuroCAAS through IAEs.

| NEUROCAAS IAEs |  |  |  |
| --- | --- | --- | --- |
| Analysis Name | Paper | Subfield | Description |
| CaImAn | (Giovannucci et al., 2019) | Calcium Imaging | Integrated python toolbox for large scale Calcium Imaging data Analysis and behavioral analysis. |
| DeepLabCut | (Mathis et al., 2018) | Pose Tracking | Markerless pose estimation of user-defined features with deep learning for all animals, including humans. |
| PMD | (Buchanan et al., 2018) | Functional Imaging | Penalized Matrix Decomposition for Denoising and Compression of Functional Imaging Data. |
| LocaNMF | (Saxena et al., 2020) | Widefield Imaging | Region-based Decomposition for Widefield Calcium Imaging Data. |
| YASS | (Lee et al., 2017) | Spike Sorting | YASS is a spike sorting pipeline developed for high-firing rate, high-collision rate retinal recordings. |
| Latent Factor Analysis for Dynamical Systems (LFADS) | (Sussillo et al., 2016) | Probabilistic inference on time series | Deep learning method to infer latent dynamics from single-trial neural spiking data. |
| DeepGraphPose | (Wu et al., 2020) | Pose Tracking | DGP is a semi supervised model which can run on top of other tracking algorithms, such as DLC. |
| BehaveNet / Partitioned-subspace VAE | (Batty et al., 2019)<br>(Whiteway et al., 2021) | Behavioral Video Analysis | Nonlinear embedding and Bayesian neural decoding of behavioral videos. |
| BarDensr | (Chen et al., 2020) | Spatial Transcriptomic Imaging | BarDensr (BARcode DEMixing through Non-negative Spatial Regression) for demixing spatial transcriptomic imaging data. |
| Kalman Filter/Smoother (Linear Dynamical System) | (Minka, 1999) | Probabilistic inference on time series | Time-invariant model for tracking a single object in a continuous state space. |
| Emergent Property Inference | (Bittner et al., 2019) | Likelihood free inference | A method for learning parameter distributions on models constrained to produce certain desirable phenomena in their output. |
| DLC Tracking (Polleux) | N/A | Pose Tracking | Markerless pose estimation with postprocessing to quantify freezing behavior. |
| Ensemble DGP | N/A | Pose Tracking | Ensemble DGP independently trains N different DeepGraphPose models that differ only in the sequence of minibatches that they see, and generates a consensus prediction from them. |

#### 2.1.2 Job managers for system infrastructure

Although many parts of system level infrastructure can be summarized in the IAE or the resource bank (discussed next), one crucial aspect that must be treated independently is setup of a communication channel between the user, who provides analysis inputs, and the NeuroCAAS infrastructure stack that processes these inputs. This setup is the responsibility of the NeuroCAAS job manager, which associates user submitted datasets to the appropriate IAEs on the right hardware, writes status messages back to the user, and monitors analysis progress, similar to a cluster workload manager like slurm (Yoo et al., 2003) (Figure 2, blue arrows). However unlike cluster managers, the NeuroCAAS job manager does not assign jobs to running infrastructure, but rather activates and de-activates the relevant infrastructure stack at the beginning and end of the job, circumventing the need for manual intervention to manage any part of the infrastructure stack (Figure 2; black arrows). The job manager for each analysis is specified with a “protocol” in code that allow it to to configure and take apart infrastructure programatically. Job manager protocols can manage complex behavior, handling large scale analyses that span multiple hardware instances, as we will demonstrate in our NeuroCAAS native analyses (see Figure 8).

#### 2.1.3 Resource banks for hardware infrastructure

Perhaps the most unintuitive infrastructure component to specify in code is computing hardware. The requirement that specific computing hardware be allocated on demand from code is handled by NeuroCAAS’s resource bank. The NeuroCAAS resource bank can make hardware available through individual *instances*: bundled collections of virtual cpus, memory, and gpus, similar to a compute node on an on-premises cluster. However unlike an on-premise computing cluster, the NeuroCAAS resource bank is built upon globally available, virtualized compute hardware offered through the public cloud (currently Amazon Web Services). At any time, the resource bank can draw on an effectively limitless volume of compute resources to emulate a wide range of familiar hardware instances (e.g. personal laptop, on premise cluster) in order to execute a particular task (Figure 2, bottom right). We have configured the resource bank with security and privacy features to ensure that hardware can only be accessed by the appropriate job manager, and individual instances generate diagnostics that can be used to monitor for aberrant behavior, providing easy management at the scale provided on this platform. Furthermore, hardware can be provisioned in a dataset-dependent manner, adjusting the size of storage volumes, memory, or computing resources as needed (see §5.1 for more details). Each configuration of a hardware instance is named with a unique identifier, allowing identical hardware to be activated at will. Fundamentally, the resource bank allows us to identify the hardware lifecycle with the lifecycle of a particular analysis job, providing efficient, custom built compute to each processing job and eliminating any upfront costs to users or developers before trying out/developing a new analysis (Figure 4B).

#### 2.1.4 Blueprints for instant reproducibility

For any given analysis, each of NeuroCAAS’s infrastructure components has a specification in code (IAEs have IDs, job managers have protocols, and the resource bank registers identifiers for specific hardware instances). The collection of all infrastructure identifying code associated with a given NeuroCAAS analysis is stored in the blueprint of that analysis (Figures 2, 5, top right), from which new instances of the infrastructure stack can be deployed at will, providing reproducibility by design. Despite sustained efforts to promote reproducible research, Buckheit and Donoho (1995), in many typical cases data analysis remains frustratingly non-reproducible (Crook et al., 2013, Gorgolewski et al., 2017, Stodden et al., 2018, Raff, 2019). NeuroCAAS sidesteps all of the typical barriers to reproducible research by tightly coupling the creation of infrastructure stacks to their documentation. Paired with a simple version control system, blueprints provide instant easy reproducibility across the entire development history of any data analysis, to anyone using NeuroCAAS.

**Figure 5:**
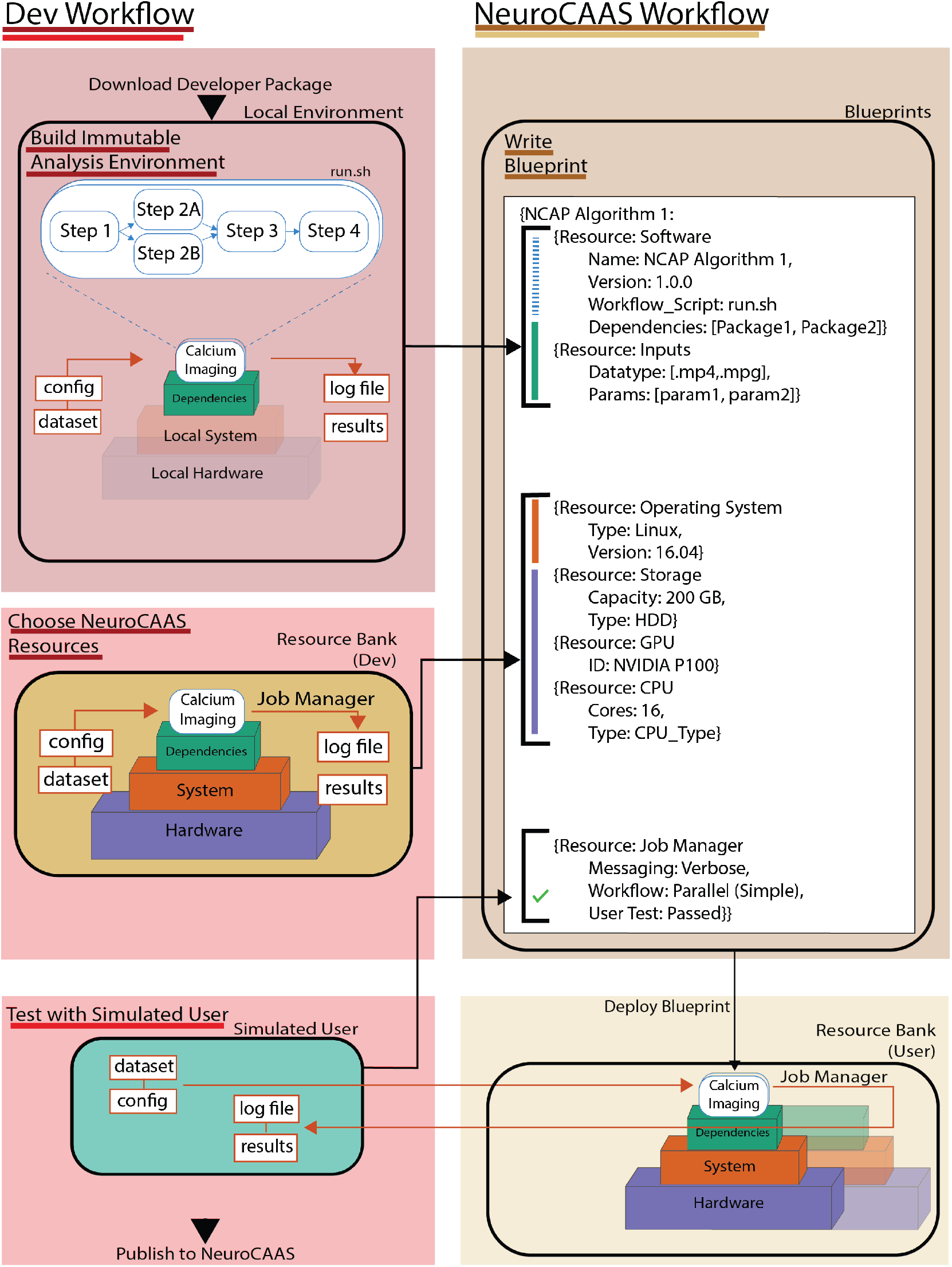
Overview of NeuroCAAS Developer Workflow. Left indicates the developer’s experience; right indicates the work that NeuroCAAS performs. The developer begins by downloading the developer package https://github.com/cunningham-lab/neurocaas_contrib and building an immutable analysis environment script on their local machine. After determining workflow and optionally installing analysis software into an IAE, the developer locally tests that sample data and config files yield expected logs and results. Once satisfied, the developer updates a blueprint with IAE specifications. Next, developers configure system and hardware settings by setting up their IAE, complete with sample data and parameters, on the NeuroCAAS resource bank. Configuration and updates to an IAE are done identically to initial local build, allowing for IAEs custom-built for powerful cloud resources. Finally, developers simulate NeuroCAAS user accounts and trigger analyses with their blueprint to ensure that the blueprint they have written function as intended end-to-end before publishing their blueprint for use.

### 2.2 Using and Developing Analyses on NeuroCAAS

NeuroCAAS supports any interface that allows users to transfer data files to and from the cloud. The standard interface to NeuroCAAS is a website, www.neurocaas.org, where users can sign up for an account, browse core analyses, deposit data and monitor analysis progress until results are returned to them. The cost of using these analyses is directly proportional to analysis duration and the type of cloud resources used to construct the relevant infrastructure stack (Figure 4B).

For comparison, Figure 4 qualitatively illustrates the accumulating inefficiencies of time and cost in every IaGS pipeline (foreshadowing the cost efficiency of NeuroCAAS’s approach, detailed in §3.3). IaGS begins with a number of time-consuming manual steps, including hardware acquisition, hardware setup, and software installation (Figure 4A). With a functional infrastructure stack in hand, the user must prepare datasets for analysis, manually recording analysis parameters and monitoring the system for errors as they work. While parallel processing is possible, it must be scripted by the user, and in many cases datasets are run serially. What results from IaGS is massive inefficiency of time and resources. Users must also support the cost of new hardware “up front,” before ever seeing the scientific value of the infrastructure that they are purchasing. Likewise, labs or institutions must pay support costs to maintain infrastructure when it is not being used, and replace components when they fail or become obsolete (Figure 4B). Two editorial remarks bear mentioning at this point: first, the stark difference laid out in Figure 4 is the essence of IaGS vs IaC, and explains the dominance of IaC in modern industrial settings. Second, NeuroCAAS is and will remain an open-source tool for the scientific community, in keeping with its sole purpose of improving the reproducibility and dissemination of neuroscience analysis tools.

To make NeuroCAAS accessible to developers, we built a suite of developer tools that streamlined the process of migrating an existing analysis to the NeuroCAAS web service, available as a python package. These developer tools abstract away the cloud infrastructure that NeuroCAAS is built on, allowing for analysis development entirely from the command line. In brief, the development process can be summarized as incrementally filling in the blueprint of a new analysis (Figure 5). First, developers configure existing analysis code (e.g. a Github repo) with scripts to be triggered by data and parameter input and stored in an immutable analysis environment (Figure 5, top left). Next, developers can interactively configure their immutable analysis environment with hardware available in the resource bank (Figure 5, middle). Finally, developers can simulate user accounts, and test the process of submitting data and awaiting results in an end-to-end manner using the job manager (Figure 5, bottom). See https://neurocaas.readthedocs.io/en/latest/index.html for a detailed developer guide.

#### 2.2.1 Testing the NeuroCAAS usage model

Over a period of 10 months, we opened the NeuroCAAS platform to a group of alpha testers (users and developers), and analyzed their development and usage patterns to optimize the design of NeuroCAAS. We recorded the number, duration, and parallelism (number of individual datasets analyzed) of jobs launched by users, and collected the results in Figure 6. This graph suggests the co-occurrence of different usage patterns: a large number of single dataset jobs are suggestive of one-off exploratory use, while there is also a considerable proportion of jobs that leverage parallelism, running analyses on anywhere from 2 to 70 datasets at a single time. These results motivated us to offer unlimited parallelism as part of the NeuroCAAS platform by default (see Figure 8A), as long as users adhered to a pre-determined cloud compute budget (initially fixed at $300 per user).

**Figure 6:**
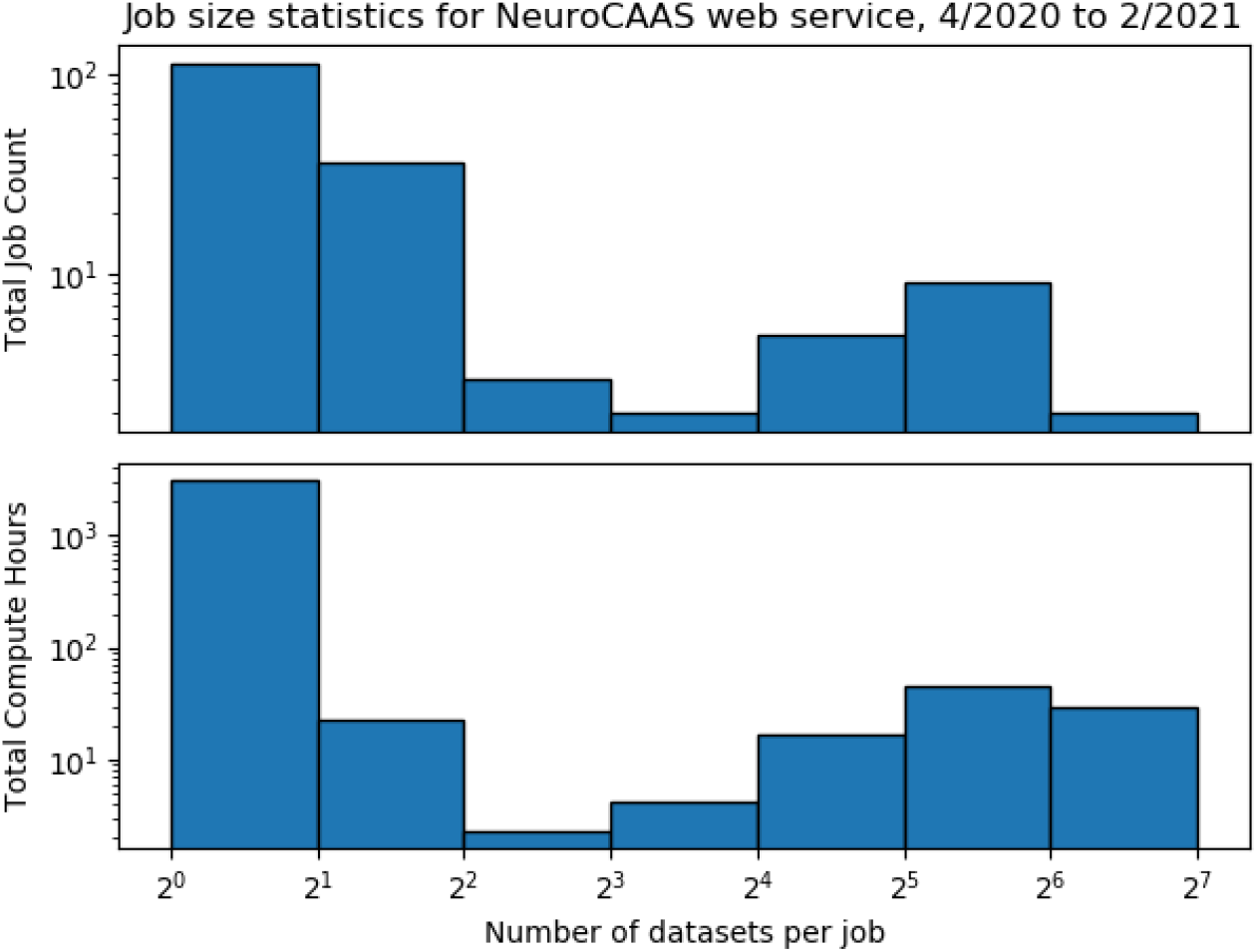
Usage of NeuroCAAS Platform. (Top) Aggregated job count across all analyses offered on the NeuroCAAS web service as a function of job parallelism-i.e. number of individual IAEs and hardware instances deployed per job. (Bottom) Corresponding aggregate compute hours devoted to analysis as a function of job parallelism.

During this test period, we also observed that NeuroCAAS benefits collaborative work between experimentalists and analysis developers. If users found bugs in the course of running analyses, this bug was very easily communicated to developers, as the only variables needed to reproduce the bug are the dataset and configuration file run by the user (without needing to know details of the user’s local operating system or software environment). In response, the developer modified the analysis blueprint accordingly, solving the issue for all future use. The replicability of errors is a major issue in open source software development, and we found that blueprint based error correction significantly eased the burden of supporting analyses on developers. In several cases, individual users also moved from exploring the generic analyses we made available on NeuroCAAS to customized versions with tailor-made pre and post processing routines, which were quickly developed by copying and modifying existing blueprints. Many of these can be found as “custom analyses” on the NeuroCAAS website.

### 2.3 Existing infrastructure tools and platforms

Figure 1 illustrates that the infrastructure stack for any analysis pipeline has many codependent parts. Existing approaches to infrastructure in neuroscience consist of platforms that group together some subset of infrastructure components and package together an existing set of analyses that rely on this shared infrastructure. Although we do not attempt an exhaustive review here, these platforms generally focus on a particular subfield of neuroscience data analysis tools (cell segmentation, human brain imaging, genomics, image processing). In Figure 7, we plot a variety of popular neuroscience analyses onto a space defined by 1) their place in the adoption lifecycle and 2) corresponding infrastructure needs. We overlay several exemplar platforms, covering those portions of analysis space that they support. The degree to which a platform’s support extends to the right defines its *accessibility,* or the ease with which developers can set up/update their analyses on the platform, and the ease with which users can begin to process data with them. Accessibility is especially important for analyses that are still early in the adoption lifecycle with active development and a growing user base. Likewise, the degree to which a platform’s support extends upwards defines its *scale*, a one dimensional approximation of the infrastructure needs for which it can provide. While the exact positioning of these analyses and platforms is subjective and dynamic, there are general characteristic features of the analysis platform landscape that we discuss in what follows.

**Figure 7:**
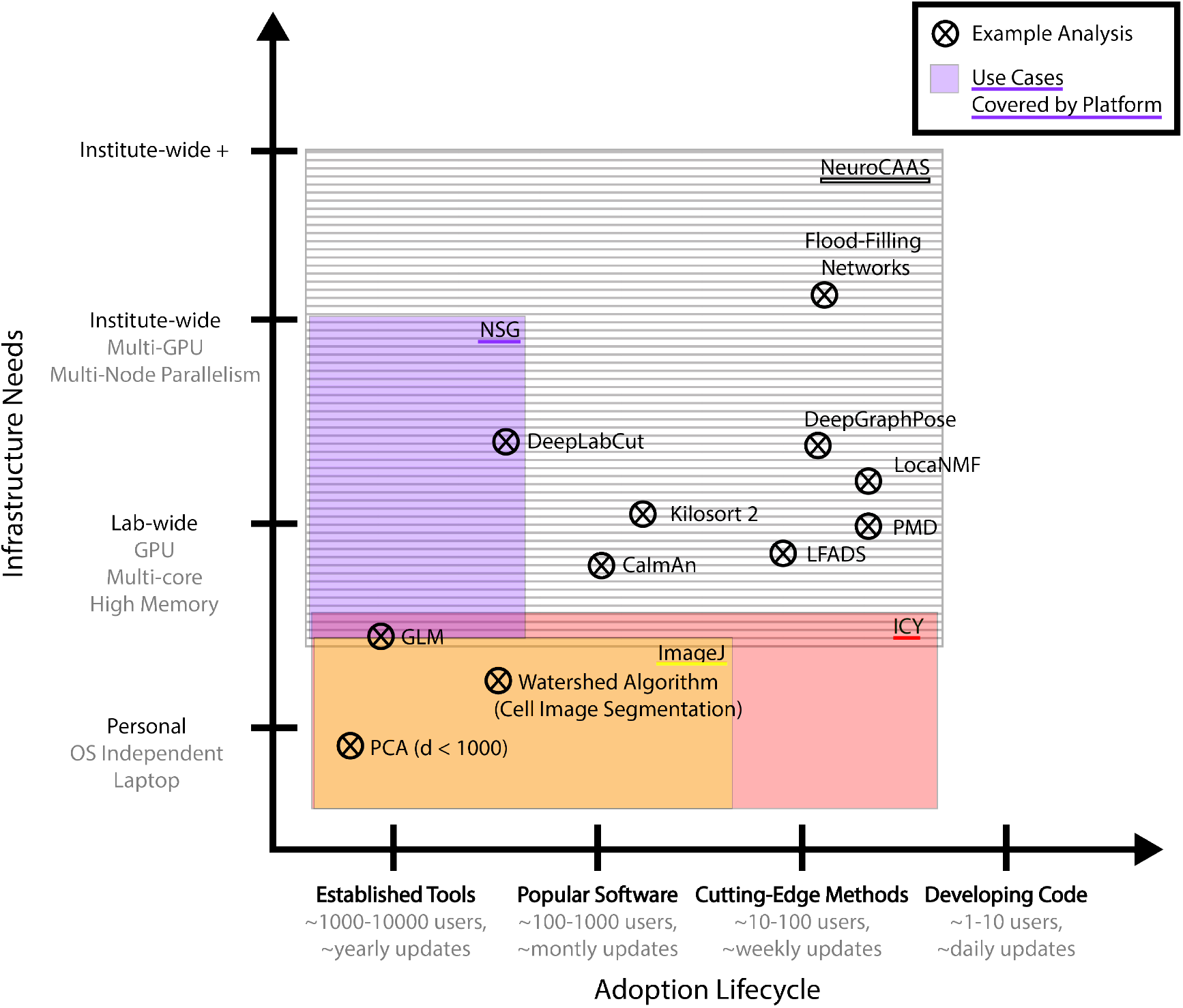
Landscape of cellular/circuit-level neuroscience analysis platforms. We organize a variety of popular analyses (denoted by crosses) in terms of their place in the adoption lifecycle (number of users, rate of software updates), and their infrastructure needs. We overlay a representative collection of analysis platforms, indicating the parts of analysis space that are covered by a given platform. Notably, there is a significant gap in the existing landscape that NeuroCAAS addresses with an IaC approach. (Example analyses: (Goodman and Brette, 2009, Pnevmatikakis et al., 2016, Mathis et al., 2018, Pachitariu et al., 2016, Pandarinath et al., 2018, Januszewski et al., 2018, Saxena et al., 2020, Buchanan et al., 2018, Graving et al., 2019); representative platforms: (Sanielevici et al., 2018, Chaumont et al., 2012, Schneider et al., 2012).

Platforms like CellProfiler (Carpenter et al., 2006), Ilastik (Sommer et al., 2011) (cell-based image processing), Icy (Chaumont et al., 2012), ImageJ (Schneider et al., 2012) (generic bioimage analysis), BIDS Apps (Gorgolewski et al., 2017) (MRI analyses for Brain Imaging Data Structure format), and Bioconductor (Amezquita et al., 2019) (genomics) have all achieved success in the field by packaging together well developed analyses with necessary software dependencies, and system management tools offered through intuitive streamlined user interfaces. These platforms can all be downloaded and installed on a user’s local infrastructure. Most of these local platforms also have an open contribution system for interested developers to build or extend custom analyses. Local platforms are thus highly accessible to both developers and users, but are in the large majority of cases installed on local hardware, limiting their scale (Figure 7, bottom).

In contrast, platforms like the Neuroscience Gateway (NSG) (Sanielevici et al., 2018) (specializing in neural simulators), Flywheel (`flywheel.io`) (emphasizing fMRI and medical imaging), and neuroscience-focused research computing clusters host computation remotely, allowing them to offer hardware at scale (through the XSEDE [Extreme Science and Engineering Discovery Environment] portal (Towns et al., 2014), the public cloud and on-premises hardware, respectively). These remote platforms offer powerful compute, but at the cost of accessibility to users, who must adapt their software and workflow to new conventions (i.e. wait times for jobs to run on shared resources, prepackaged coding environments, limitations on concurrency) in order to make use of offered hardware. As a particular example, NSG requires users to submit a script that they would like to have run on a compute node in the language of their chosen analysis, making it more similar to a traditional on-premises cluster in usage than NeuroCAAS. NSG also restricts jobs to run on one compute instance at a time, making it incompatible with the usage model and novel analyses we will present. Finally, NSG does not have an open system for contributing new analyses, making it more suitable for established analysis tools with stable, indepenently maintained documentation. Likewise, Flywheel (Flywheel Exchange, 2020) (with a focus on human brain imaging tools), requires a paid subscription in order to access its services for both analysis use or development, making it more suitable to support specific groups and collaborations with a clear interest in using or developing a shared set of analysis tools, rather than the general purpose infrastructure goals of NeuroCAAS. These platforms are best for committed, experienced users who already work with the analyses available on the remote platform. It is also more difficult to contribute new analyses to these platforms than their locally hosted counterparts (see Figure 7, left side). This difficulty makes them less suitable for actively developing or novel analyses, as updates may be slow to be incorporated, or introduce breaking changes to user written scripts.

Although undeniably useful, local and remote platforms operate on a tradeoff that forces researchers to choose between accessibility and scale. While these platforms often concentrate on applications that mitigate the effects of this tradeoff, there are many popular analyses that would not be suitable for existing analysis platforms (see Figure 7, center). Furthermore, with both types of platforms, users are still required to work directly with analysis infrastructure, whether by installing new tools onto one’s personal infrastructure or preparing code and dependencies to run in a remote (and sometimes variably allocated) infrastructure stack. Both situations can introduce problematic IaGS issues and impact the reproducibility of derived results.

Beyond platforms that explicitly offer analysis infrastructure, there are tools that make given infrastructure more reproducible, offering automated documentation and logging tools (Greff et al., 2017), tracing the system call of an analysis run (Chirigati et al., 2016), or aggregating data analyses onto an open source operating system/-Docker container (Halchenko and Hanke, 2012). Though admirable, these projects all require a non-trivial effort by users and developers to learn and integrate a new tool into their work. Furthermore, the evolving landscape of new analysis tools and platforms makes it difficult to develop standalone reproducibility tools that can cover the variety of analyses and infrastructure stacks used in our field.

## 3 Results

NeuroCAAS’s comprehensive approach to analysis infrastructure solves issues that IaGS based approaches can not. Here, we demonstrate how IaC enables simple, powerful infrastructure design for the analysis of large datasets (§3.1) and for the deployment of large-scale deep learning pipelines (§3.2), and show that the cost and accessibility of NeuroCAAS offers distinctive benefits to popular analyses across the tested range of use cases (§3.3).

### 3.1 IaC for Large Data: Protocol for Widefield Imaging

Some of the most infrastructure-intensive analyses in neuroscience are preprocessing techniques that work directly on big data. The high degree of automation required to work with large datasets demands many separate preprocessing steps, creating the need for unwieldy multi-analysis infrastructure stacks- individual stacks that support the infrastructure needs of multiple core analyses at the same time. A notable example is data generated by wide-field imaging of calcium indicators (WFCI)- a high-throughput imaging technique that collects activity dependent fluorescence signals across the entire dorsal cortex of an awake, behaving mouse (Couto et al., 2020), potentially generating terabytes of data across chronic experiments. The protocol paper Couto et al. (2020) describes a complete WFCI analysis that links together cutting edge data compression/denoising with demixing techniques designed explicitly for WFCI (via Penalized Matrix Decomposition, or PMD (Buchanan et al., 2018) and Lo-caNMF (Saxena et al., 2020), respectively). Each of these analyses depends upon its own specialized hardware and requires operating system dependent installation, creating many competing requirements on a multi-analysis infrastructure stack that are difficult to satisfy in practice. While we offer a NeuroCAAS implementation of the described WFCI analysis in Couto et al. (2020), we do not discuss how NeuroCAAS addresses the issue of multi-analysis infrastructure stacks, which can pose IaGS challenges even to our blueprint based infrastructure.

Instead of working with multi-analysis infrastructure stacks, NeuroCAAS introduces a new way to design analyses that span multiple independent steps. On NeuroCAAS, existing analysis components stored in independent blueprints can be linked together through their inputs and outputs (Figure 8B). We employ this design to build a complete analysis for WFCI, where the initial steps of motion correction, denoising, and hemodynamic correction of the data are performed on a stack that emphasized multicore parallelism (64 cores) to suit the matrix decomposition algorithms employed by PMD. Upon termination of this first step, analysis results are not only returned to the user, but also used as inputs to a create a second stack, performing demixing with LocaNMF on infrastructure supporting a high performance GPU. This modular organization improves the performance and efficiency of each analysis component (see Figure 10), and also allows users to run steps individually if desired, giving them the freedom to interleave existing analysis pipelines with the components offered here. This complete WFCI analysis can be controlled through a custom built graphical user interface (GUI), where the user can interactively initialize data processing, submit jobs to NeuroCAAS, and explore analysis results, providing a high degree of accessibility for this complex, multi-step pipeline. Importantly, NeuroCAAS balances this accessibility with scale, as the performance of our WFCI analysis does not depend on the infrastructure available to the user. For example, users can simultaneously launch many analyses and have them run in parallel through the GUI, easily conducting a formal, well documented hyperparameter search across all parts of their analysis simultaneously. Researchers can find the GUI for this WFCI analysis with NeuroCAAS integration at https://github.com/jcouto/wfield.

**Figure 8:**
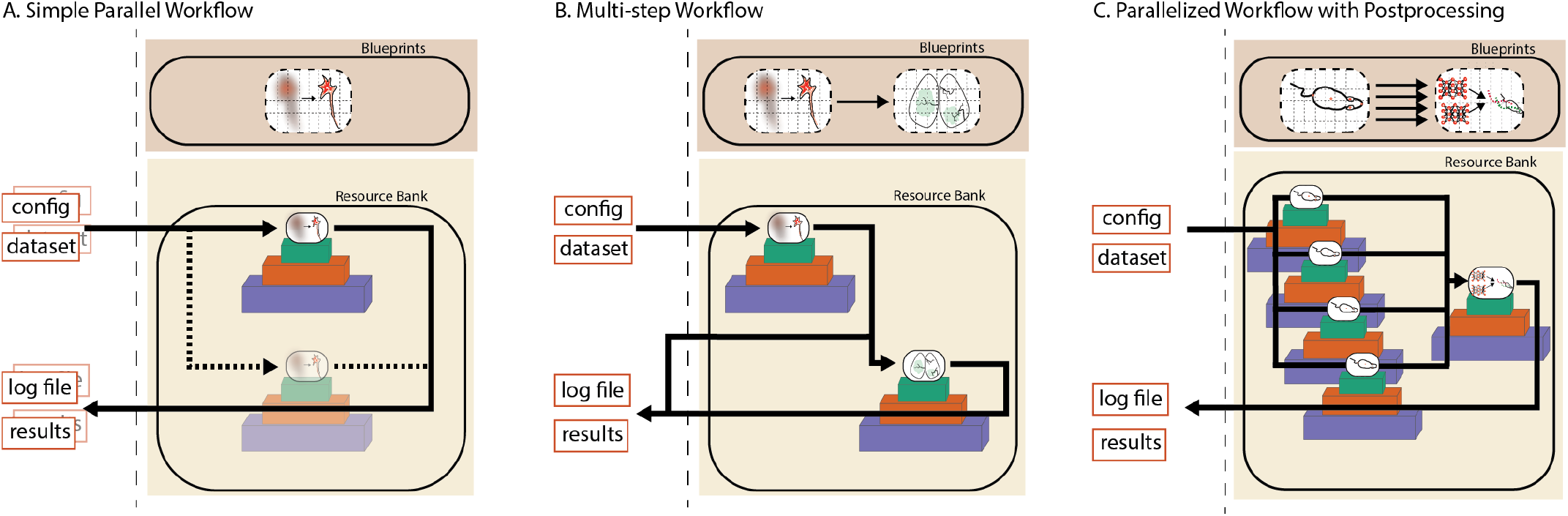
NeuroCAAS supports multi-stack design patterns. NeuroCAAS allows analyses to easily span multiple infrastructures as is most appropriate for a given task with flexible job manager protocols. A. Default workflow on NeuroCAAS specifies that infrastructure should be created on demand when datasets and analyses are submitted. If more than one dataset is submitted, we automatically create separate infrastructure for each. B. We can also specify in blueprints that the output of one analysis should feed input to another. With this chained structure, multiple analysis components with different infrastructure needs are seamlessly combined on demand. Intermediate results are returned to the user so that they can be examined and visualized as well. This is the job structure for widefield imaging analysis, §3.1. C. The easy parallelism of panel A and the chained structure of panel B can be combined in a single workflow to support batch processing pipelines with a separate postprocessing step. This is the job structure for ensemble markerless tracking, §3.2.

Additionally, for developers the separation of different analysis components into different blueprints vastly simplifies the effort required to combine different cutting edge analysis components. Existing domain specific projects such as CaImAn (Giovannucci et al., 2019) for cellular resolution calcium imaging or SpikeInterface (Buccino et al., 2020) for electrophysiology data demonstrate that making big data analyses compatible with standard hardware can require significant revision of the original analysis implementation. In contrast, our WFCI analysis is made directly available to users on powerful remote hardware without the need to revise existing analysis components, and can easily be extended to subsequent processing steps. To our knowledge, there is no other system with the accessibility and scale to link together cutting edge analyses across separate infrastructures, and make them available directly to the research community.

### 3.2 IaC for Deep Learning Models: Ensemble Markerless Tracking

The black box nature of deep learning can generate sparse, difficult to detect errors that reduce the benefits of deep learning based tools for sensitive applications. For modern markerless tracking analyses built on deep neural networks (Mathis et al., 2018, Graving et al., 2019, Nilsson et al., 2020, Wu et al., 2020), these errors can manifest as “glitches” (Wu et al., 2020), where a marker point will jump to an incorrect location, often without registering as an error in the network’s generated likelihood metrics (see Figure 9).

**Figure 9:**
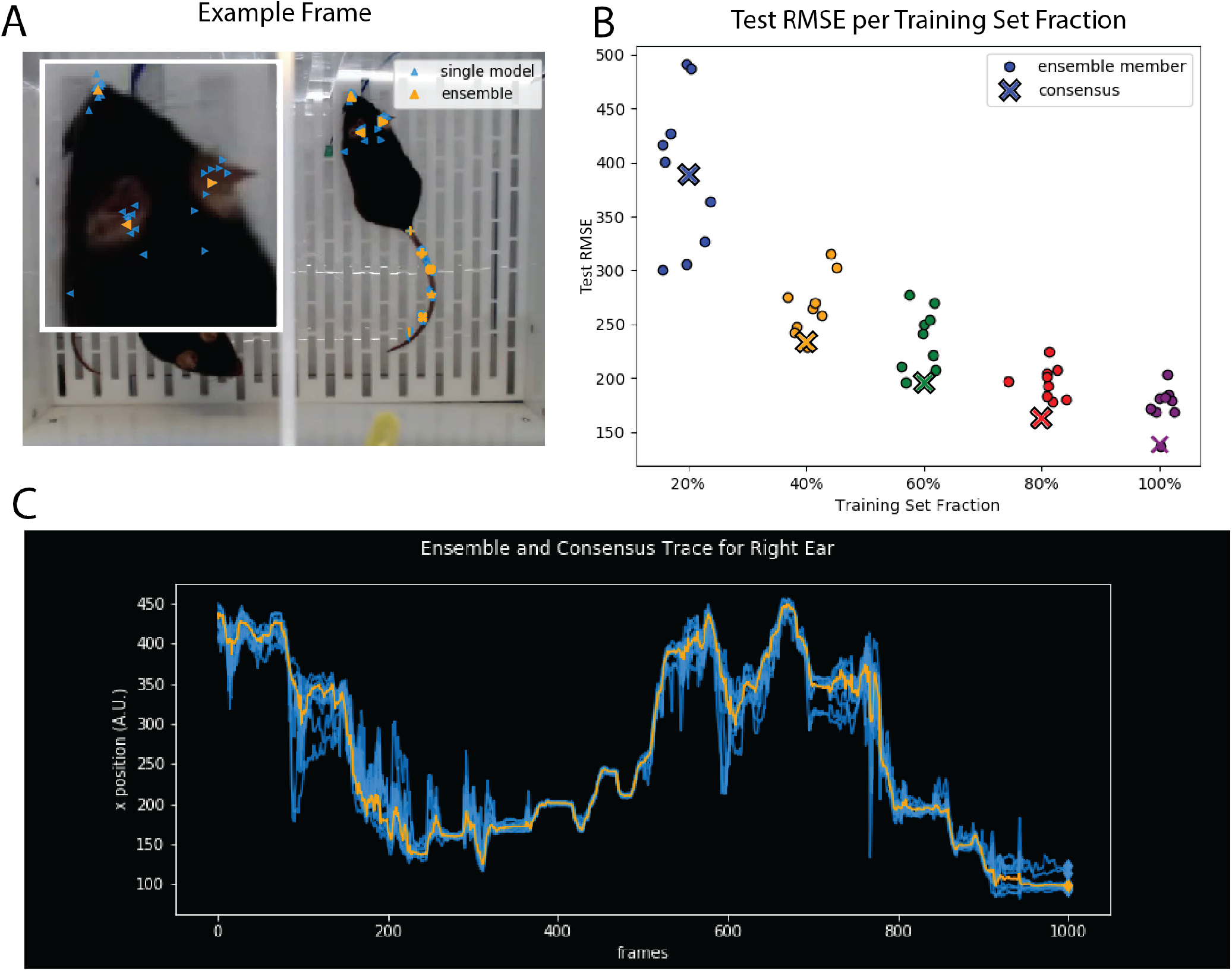
Ensemble markerless tracking. A) shows an example frame from a mouse behavior dataset (courtesy of Erica Rodriguez and C. Daniel Salzman) tracking keypoints on the top down view of a mouse, as analyzed in Wu et al. (2020). Each marker shape corresponds to a different body part, with blue markers representing the output of individual tracking models, and orange markers representing the consensus. Inset image shows tracking performance on the nose and ears of the mouse. Across body parts, certain networks in the ensemble converge to a solution that causes them to incorrectly localize keypoints (see inset, bottom left), while the aggregate detection (labeled consensus) stays close to the true body part locations. B) shows consensus test performance vs. test performance of individual networks on a dataset with ground truth labels. Performance on the test set improves as a function of training set size, but individual networks can behave very differently, even trained on the same data. In comparison, the consensus detection reliably finds solutions close to the optimum for a given training set size. C) shows traces from 9 networks (blue) + consensus (orange). Traces from individual networks diverge on challenging portions of the video, while consensus tracking remains smooth due to aggregation of detections from individual networks. Across the entire figure, ensemble size = 9. A and C correspond to traces taken from the 100% split in B with 20 training frames.

One general purpose approach to combat the unreliable nature of individual machine learning models is *ensembling* (Dietterich, 2000): instead of working with a single model, a researcher simultaneously prepares multiple models on the same task, subsequently aggregating their outputs into a single consensus output. Ensemble methods have been shown to be effective for deep networks in a variety of contexts, (Lakshminarayanan et al., 2016, Fort et al., 2019, Ovadia et al., 2019), but they confer a massive infrastructure burden if run on limited local compute resources: researchers must simultaneously train, manage, and aggregate outputs across many different deep learning models, incurring either prohibitively large commitments to deep learning specific infrastructure and/or infeasibly long wait times.

In contrast, NeuroCAAS enables easy and routine deployment of ensemble methods. Using NeuroCAAS, we designed an analysis which takes input training data, and distributes it to *N* different instances of the same base tracking model (Figure 8C). For the application shown here, we used DeepGraphPose (Wu et al., 2020) as our base tracking model; the N instances differ only in the minibatch order used for training. The results from each trained model are then used to produce a consensus tracking output, taking each individual model’s estimate of part location across the entire image (i.e. the confidence map output) and averaging these estimates. Even with this relatively simple approach, we find the consensus tracking output is robust to the errors made by individual models (Figure 9A,C). Furthermore, consensus performance is maintained even when we significantly reduce the size of the training set (Figure 9B). Finally, in Figure 9C, we can see that there are portions of the dataset where the individual model detections fluctuate around the consensus detection. This fluctuation offers an empirical readout of tracking difficulty within any given dataset; frames with large diversity in the ensemble outputs are good candidates for further labeling, and could be easily incorporated in an active learning loop. Overall, Figure 9 shows that with the scale of infrastructure available on NeuroCAAS, ensembling can easily improve the robustness of markerless tracking, naturally complementing the infrastructure reproducibility provided by the platform.

NeuroCAAS is uniquely capable of providing the flexible infrastructure necessary to support a generally available, on-demand ensemble markerless tracking application. To our knowledge, none of the remote platforms with the scale to support an IaGS approach to markerless tracking (e.g. on premise clusters, Google Colab, NSG (Sanielevici et al., 2018)) can satisfactorily alleviate the burden of a deep ensembling approach, still forcing the user to accept either long wait times or tedious manual management of infrastructure. These limitations also prohibit use cases involving the quantification of ensemble behavior across different parameter settings (c.f. Figure 9B, where we trained 45 networks simultaneously).

### 3.3 NeuroCAAS is faster and cheaper than local “on-premises” processing

NeuroCAAS offers a number of major advantages over IaGS: reproducibility, accessibility, and scale, whether we compare against a personal workstation or resources allocated from a locally available cluster. However, since NeuroCAAS is based on a cloud computing architecture, one might worry that data transfer times (i.e., uploading and downloading data to and from the cloud) could potentially lead to slower overall results or that the cost of cloud compute could outweigh that of local infrastructure.

Figure 10 considers this question quantitatively, comparing NeuroCAAS to a simulated personal workstation (see §5.3 for details). For the analogous comparisons against a simulated local cluster, see Figure 15. Figure 10 presents time and cost benchmark results on four popular analyses that cover a variety of data modalities: CaImAn (Giovannucci et al., 2019) for cellular resolution calcium imaging; DeepLabCut (DLC) (Mathis et al., 2018) for markerless tracking in behavioral videos; and a two-step analysis consisting of PMD (Buchanan et al., 2018) and LocaNMF (Saxena et al., 2020) for analysis of widefield imaging data. To be (extremely) conservative, we assume local infrastructure is set up, neglecting all of the time associated with installing and maintaining software and hardware.

**Figure 10:**
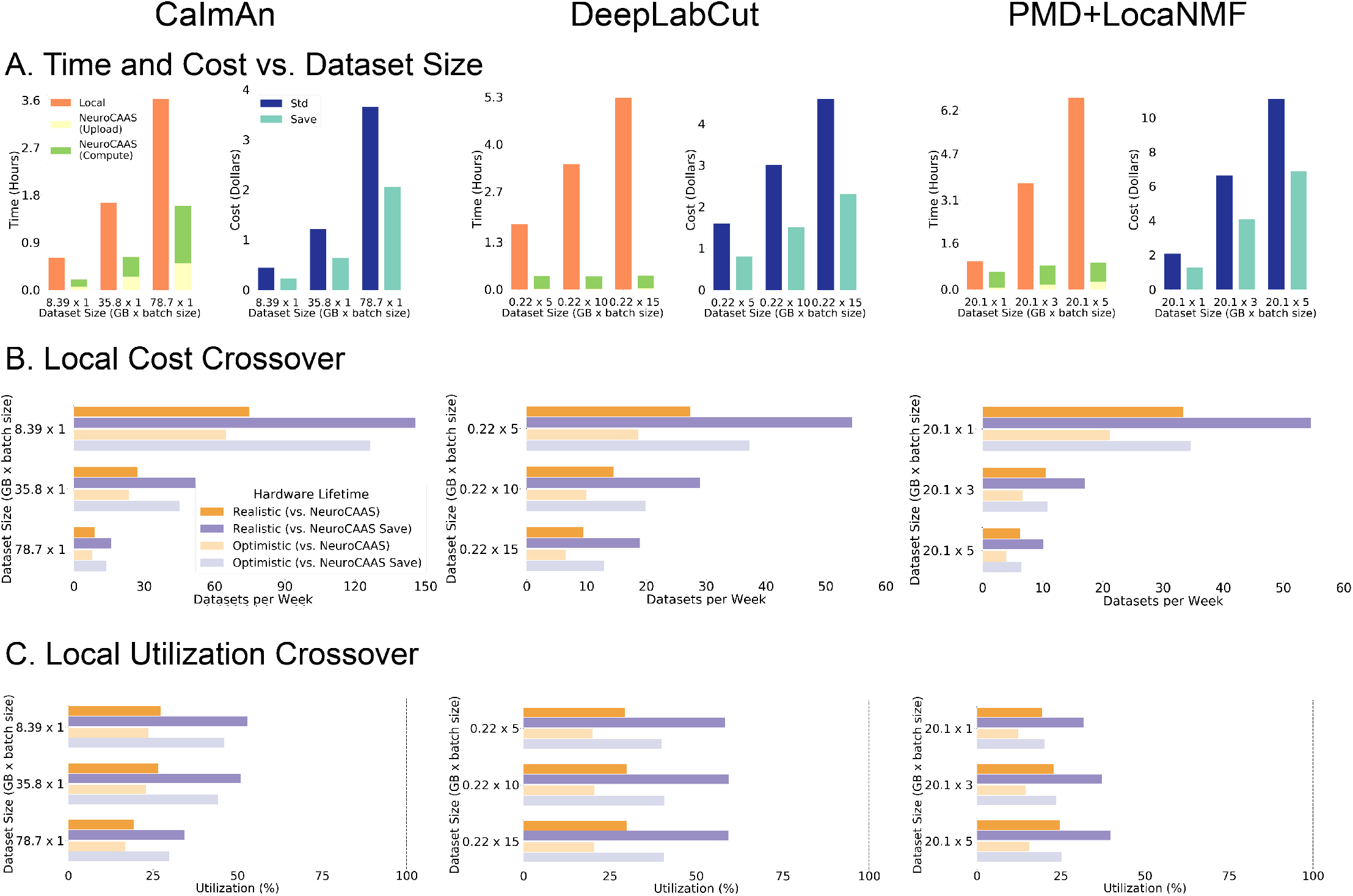
Quantitative comparison of NeuroCAAS versus local processing for three different analysis pipelines. A. Simple quantifications of NeuroCAAS performance. Left graphs compare total processing time on NeuroCAAS vs. local infrastructure (orange). NeuroCAAS processing time is broken into two parts: Upload (yellow) and Compute (green). Repeated analyses of data already in NeuroCAAS need only consider Compute times (see main text for details). Right graphs quantify cost of analyzing data on NeuroCAAS with two different pricing schemes: Standard (dark blue) or Save (light blue), offering the same analyses at a reduced price if the approximate duration of an analysis job is known beforehand. B. Cost comparison with local infrastructure. Local Cost Crossover gives the minimum per-week data analysis rate required to justify the cost of a local infrastructure compared to NeuroCAAS. We consider local pricing against both Standard and Save prices, and with Realistic (2 year) and Optimistic (4 year) lifecycle times for local hardware. C. Achieving Crossover Analysis Rates. Local Utilization Crossover gives the minimum utilization required to achieve crossover rates shown in B. Dashed vertical line indicates maximum feasible utilization rate at 100%.

Across all analyses and datasets considered in Figure 10, analyses run on NeuroCAAS were significantly faster than those run on the selected local infrastructure, even accounting for the time taken to stage data to the cloud (Figure 10A, left panes). For CaImAn, we took advantage of the fact that the algorithm was built to parallelize across multiple cores of the same machine, and chose hardware to make effective use of this implementation across data sizes (for details see Giovannucci et al. (2019), Figure 8). For DLC and PMD+LocaNMF, the NeuroCAAS compute time was effectively constant across increasing total dataset size, as we assumed data was evenly batched into subsets of approximately equal size and each batch was analyzed in its own independent instance (as in Figure 8A). These examples show that many analyses can be used efficiently on NeuroCAAS regardless of the degree to which they have been intrinsically optimized for parallelism. Finally, NeuroCAAS upload time can be ignored if analyzing data that has already been staged for processing — for example if there is a need to reprocess data with an updated algorithm or parameter setting — leading to further speedups. Next we turn to cost analyses. Over the range of algorithms and datasets considered here, we found that the overall NeuroCAAS analysis cost was on the order of a few US dollars per dataset (Figure 10A, right panels). In addition to our baseline implementation, we also offer an option to run analyses at a significantly lower price (indicated as “Std” and “Save” respectively in the cost barplots in Figure 10), if the user can upper bound the expected runtime of their analysis to anything lower than 6 hours (i.e. from previous runs of similar data, or complexity estimates).

Finally, we compare the cost of NeuroCAAS directly to the cost of purchasing local infrastructure. We use a total cost of ownership (TCO) metric (Morey and Nambiar, 2009) that includes the purchase cost of local hardware, plus reasonable maintenance costs over estimates of hardware lifetime; see §5.3 for full details. We first ask how frequently one would have to run the analyses presented in Figure 10 before it becomes worthwhile to purchase dedicated local infrastructure. This question is answered by the Local Cost Crossover (LCC): the threshold weekly rate at which a user would have to analyze data for NeuroCAAS costs to exceed the TCO of local hardware. As an example, the top two bars of Figure 10B, left, show that in order for a local machine to be cost effective for CaImAn, one must analyze ~ 100 datasets of 8.39 GB per week, every week for several years (see Table 5 for a conversion to data dimensions). In all use cases, the LCC rates in Figure 10B show that a researcher would have to consistently analyze ~ 10 — 100 datasets per week for several years before it becomes cost effective to use local infrastructure. While such use cases are certainly feasible, managing these use cases on local infrastructure via IaGS would involve an incredible amount of human labor.

In Figure 10C, we characterize this labor cost via the Local Utilization Crossover (LUC): the actual time cost of analyzing data on a local machine at the corresponding LCC rate. Across the analyses that we considered, local infrastructure would have to be dedicated to the indicated analysis for 25-50% of the infrastructure’s total lifetime (i.e. ~6-12 hours per day, every day) to achieve its corresponding LCC threshold, requiring an inordinate amount of work on the part of the researcher to manually run datasets, monitor analysis progress for errors, or build the computing infrastructure required to automate this process– in essence forcing researchers to perform by hand the large scale infrastructure management that NeuroCAAS achieves automatically. These calculations demonstrate that even without considering all of the IaGS issues that our solution avoids, it is difficult to use local infrastructure more efficiently than NeuroCAAS. Given the variability of infrastructure availability, we also provide a tool for users to benchmark their available infrastructure options against NeuroCAAS (see the instructions at https://github.com/cunningham-lab/neurocaas).

### 3.4 NeuroCAAS is offered as a free service for many users

In many cases, researchers may use infrastructure available on hand to test out analyses before purchasing a dedicated infrastructure stack for their analyses. Given the low per-dataset cost and the major advantages summarized above of NeuroCAAS compared to the current IaGS status quo, we have decided to mirror this model on the NeuroCAAS platform, and subsidize a large part of NeuroCAAS usage by the community. Users do not need to set up any billing information or worry about incurring any costs when starting work on NeuroCAAS; we cover all costs up to a per-user cap (intially set at $300). This subsidization removes one final friction point that might slow adoption of NeuroCAAS, and protects NeuroCAAS as a non-commercial open-source effort. Since NeuroCAAS is relatively inexpensive, many users will not hit the cap; thus, for these users, NeuroCAAS is offered as a free service.

## 4 Discussion

NeuroCAAS integrates rigorous infrastructure practices into neural data analysis while also respecting current development and use practices. NeuroCAAS introduces automated infrastructure management methods via our immutable analysis environments (IAEs), job managers, and resource banks that can be easily encoded in a blueprint to reliably handle the increasingly large and complex infrastructure that has become characteristic of modern approaches to neuroscience. As representatives of NeuroCAAS’s potential, we present IaC based analyses that address infrastructure issues in ways that are qualitatively distinct from an IaGS approach. Finally, we show that the scientific virtues of NeuroCAAS are accompanied by increases in efficiency, reducing both the time and cost required to run neuroscience data analyses.

The fundamental choice made by NeuroCAAS is to provide analysis infrastructure with as much automation as possible. This choice naturally makes NeuroCAAS into a *service,* such that neither analysis users nor analysis developers have to introduce a new library or framework into their analysis and development practices; rather, NeuroCAAS removes the infrastructure burden entirely. Such a choice is a tradeoff worth making explicit:

— *What* NeuroCAAS *does.* NeuroCAAS provides the user an interface to the analysis method that functions exactly as intended by the developer. For the developer, NeuroCAAS provides a user-independent approach to analysis configuration that alleviates the burden of maintaining an open source project across diverse computing environments, and simultaneously frees developers to design infrastructure to be as powerful or complicated as is optimal without being constrained by accessibility concerns. These benefits to analysis users and developers will collectively tighten the feedback loop between experimental and computational neuroscientists.
— *What* NeuroCAAS *does not do.* First, NeuroCAAS does not aim to improve the scientific use of neural data analysis algorithms. For example, if a user has data that is incorrectly formatted for a particular algorithm, the same error will happen with NeuroCAAS as it would with conventional usage. However, this statement does not suggest that analyses on NeuroCAAS are a black box. All NeuroCAAS analyses are built from open source projects, and the workflow scripts used to parse datasets and config files inside an IAE are made available to all analysis users. Furthermore, our novel analyses show that there are means of comprehensively characterizing analysis performance that only become available at scale (i.e. full parameter searches over a multi-step analysis, or ensembling to generate more reliable uncertainty metrics).

Second, NeuroCAAS is not a scientific workflow management system. Workflow management systems for neuroscience such as Datajoint (Yatsenko et al., 2015) or more general tools like snakemake (Koster and Rahmann, 2012) and the Common Workflow Language (Amstutz et al., 2016) codify the sequential steps that make up a data analysis on given infrastructure, ensuring data integrity and provenance. While the design of NeuroCAAS incorporates some level of workflow management, our main goal is not to schematize analysis workflow. Instead, our goal is to organize and automate the work that must be done to make a *given* data analysis functional, efficient, and accessible. This goal is orthogonal and complementary to applications that explicitly provide tools to make rigorous the data infrastructure connecting experiment to database and data processing.

Third, NeuroCAAS is not unstructured access to cloud computational resources. The concept of IAEs should clarify this fact: NeuroCAAS serves a set of analyses that are configured to a particular specification, as established by the analysis developer. This constraint is often ideal, since the specification is in many cases established by the analysis method’s original authors. Further, it must been noted that these constraints are the source of NeuroCAAS’s benefits to accessibility, scale, and reproducibility. Without specific structure to manage the near infinite scale of resources available on the cloud, the management of resources on the cloud easily becomes susceptible to the issues of IaGS that motivated the development of NeuroCAAS to begin with (Monajemi et al., 2019). This constraint distinguishes NeuroCAAS from data analysis offerings like Google Colab, in keeping with their differing intended use cases. With time-limited access to GPU-backed jupyter notebooks, Google Colab is useful for one-off interactive jobs, but is limited in scale and reliability of available infrastructure. Colab allows access to only one GPU at a time and a timeout of 12 hours per machine assuming constant interaction, and the specifics of resource type fluctuate without notice (Google Research, 2017). Although this statistic is unpublished, in practice unmonitored GPUs can be reclaimed in as little as 30 minutes.

By virtue of its open source code and public cloud construction, NeuroCAAS will naturally continue to evolve. First, we hope to build a community of developers who will add more analysis algorithms to NeuroCAAS, with an emphasis on subfields of computational analysis that we do not yet support. We also plan to add support for real-time processing (e.g., Giovannucci et al. (2017) for calcium imaging, or Schweihoff et al., 2019, Kane et al., 2020 for closed-loop experiments, or Lopes et al. (2015) for the coordination of multiple data streams). Second, other tools have brought large-scale distributed computing to neural data analyses (Freeman, 2015, Rocklin, 2015) in ways that conform to more traditional high performance computing ideas of scalability for applications that are less easily parallelized than those presented here. Integrating more elaborate scaling into NeuroCAAS while maintaining development accessibility will be an important goal going forwards. Third, to facilitate more interactive workflows on larger datasets, we plan further integration with database systems such as Datajoint (Yatsenko et al., 2015) and data archives like DANDI (distributed archives for neurophysiology data integration) (Dandi Team, 2019). Finally, a major opportunity for future work is the integration of NeuroCAAS with modern visualization tools. We have emphasized above that immutable analysis environments on NeuroCAAS are designed with the ideal of fully automated data analyses in mind, because of the virtues that automation brings to data analyses. However, we recognize that for some of the core analyses on NeuroCAAS, and indeed most of those popular in the field, some user interaction is required to optimize results. We will aim establish a general purpose configuration path in the spirit of the user interface we have built for our widefield calcium imaging analysis, with which analysis developers can also serve an interactive user interface without sacrificing the benefits of cost efficiency, scalability, and reproducibility that distinguish NeuroCAAS in its current form.

Longer term, we hope to build a sustainable and open-source user and developer community around NeuroCAAS. We welcome suggestions for improvements from users, and new analyses as well as extensions from interested developers, with the goal of creating a sustainable community-driven resource that will enable new large-scale neural data science in the decade to come.

## 5 Materials and methods

### 5.1 NeuroCAAS architecture specifics

The software supporting the NeuroCAAS platform has been divided into three separate Github repositories. The first, https://github.com/cunningham-lab/neurocaas is the main repository that hosts the Infrastructure-as-Code implementation of NeuroCAAS. We will refer to this repository as the *source repo* throughout this section. The source repo is supported by two additional repositories: https://github.com/cunningham-lab/neurocaas_contrib hosts contribution tools to assist in the development and creation of new analyses on NeuroCAAS, and https://github.com/jjhbriggs/neurocaas_frontend hosts the website interface to NeuroCAAS. We will refer to these as the *contrib repo* and the *interface repo* respectively throughout this section. We discuss the relationship between these repositories in the following section, and in Figure 11. At the time of submission, we have released all three of these repositories (version 1.0.1 for the source and contrib repo, version 1.0.0 for the interface repo). All releases are documented on Zenodo, with DOIs 10.5281/zenodo.4885097, 10.5281/zenodo.4884713, and 10.5281/zenodo.4851187, respectively.

**Figure 11:**
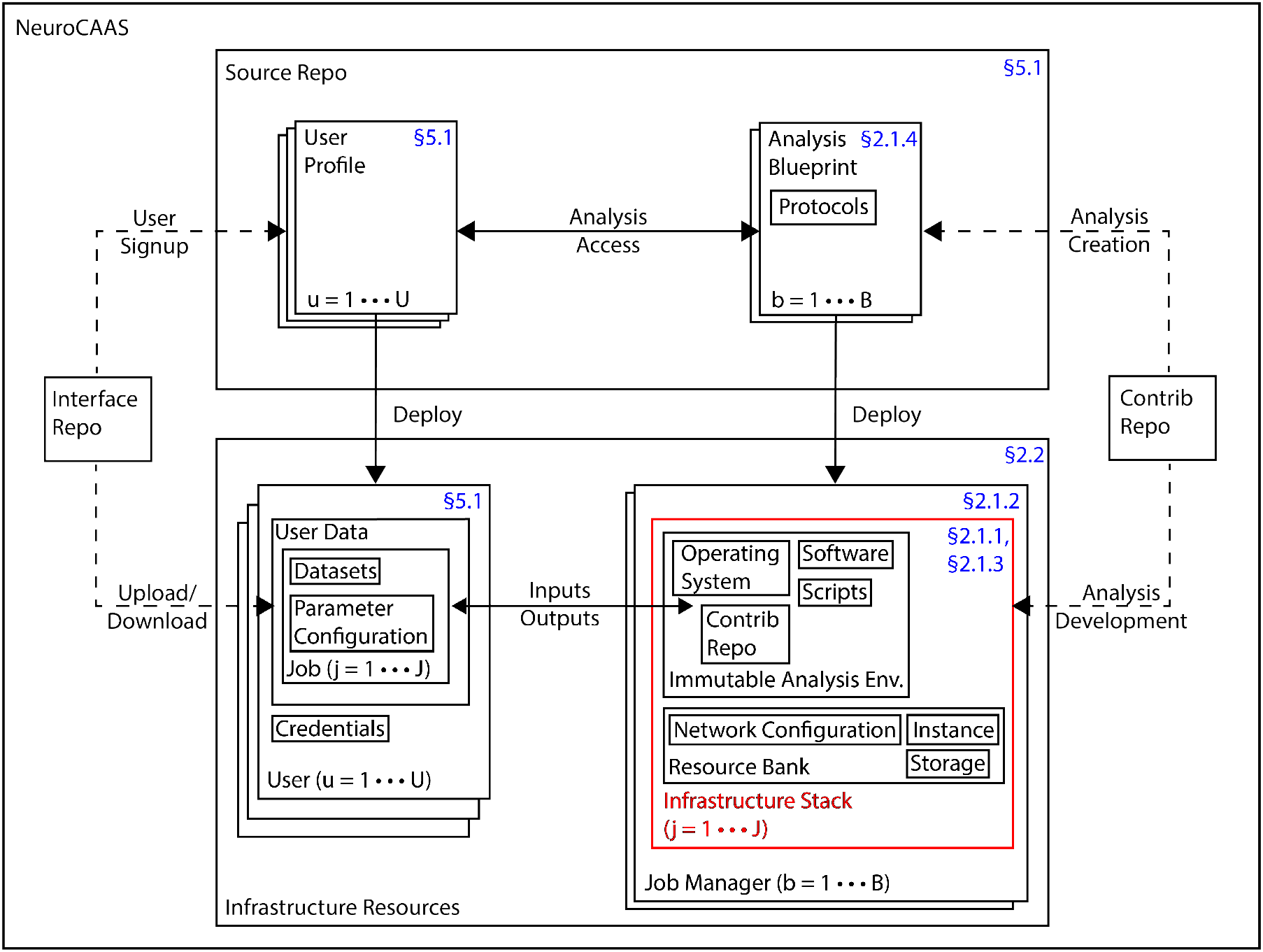
NeuroCAAS design diagram. NeuroCAAS is built with an Infrastructure-as-Code design, meaning that we first write a *source repo* (top) specifying all of the actual resources we will use to carry out data processing (bottom). The source repo (top) contains three main types of code: User Profiles, specifying relevant user data; Analysis Blueprints, describing individual analyses on NeuroCAAS, and Protocols, giving rules that describe NeuroCAAS job manager function. Each user and each analysis in NeuroCAAS has a dedicated code document, as specified by indices (u, b). All parts of the source repo can independently be *deployed,* automatically provisioning and configuring the infrastructure resources specified therein. Deployment comprehensively generates the resources necessary to run analyses on NeuroCAAS. Notably, infrastructure stacks (bottom right) are not persistent, but rather are instantiated every time users request an analysis job, specified as a combination of datasets and parameter configurations (bottom left). Job managers deploy one infrastructure stack for each requested job, as specified by the index *j*. The contrib and interface repo assist in the deployment of resources from the source repo, and and the management of resulting resources. Section numbers refer to relevant parts of the main text.

**Figure 12:**
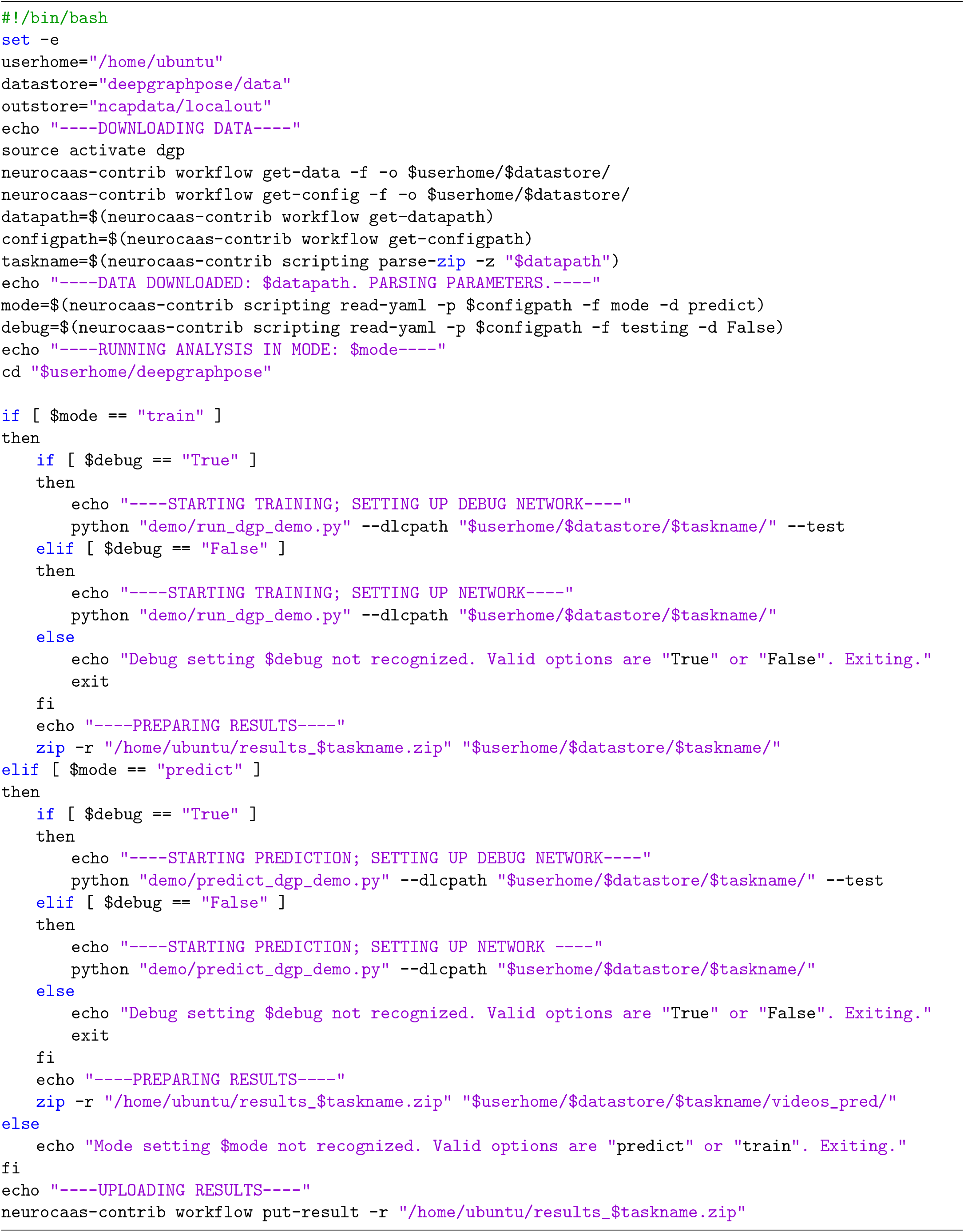
DeepGraphPose script, written in bash. Script makes heavy use of neurocaas developer tools to move data to and from NeuroCAAS data storage; see developer guide for details. Script demo/predict’dgp’demo.py has been adapted to work for any model folder.

**Figure 13:**
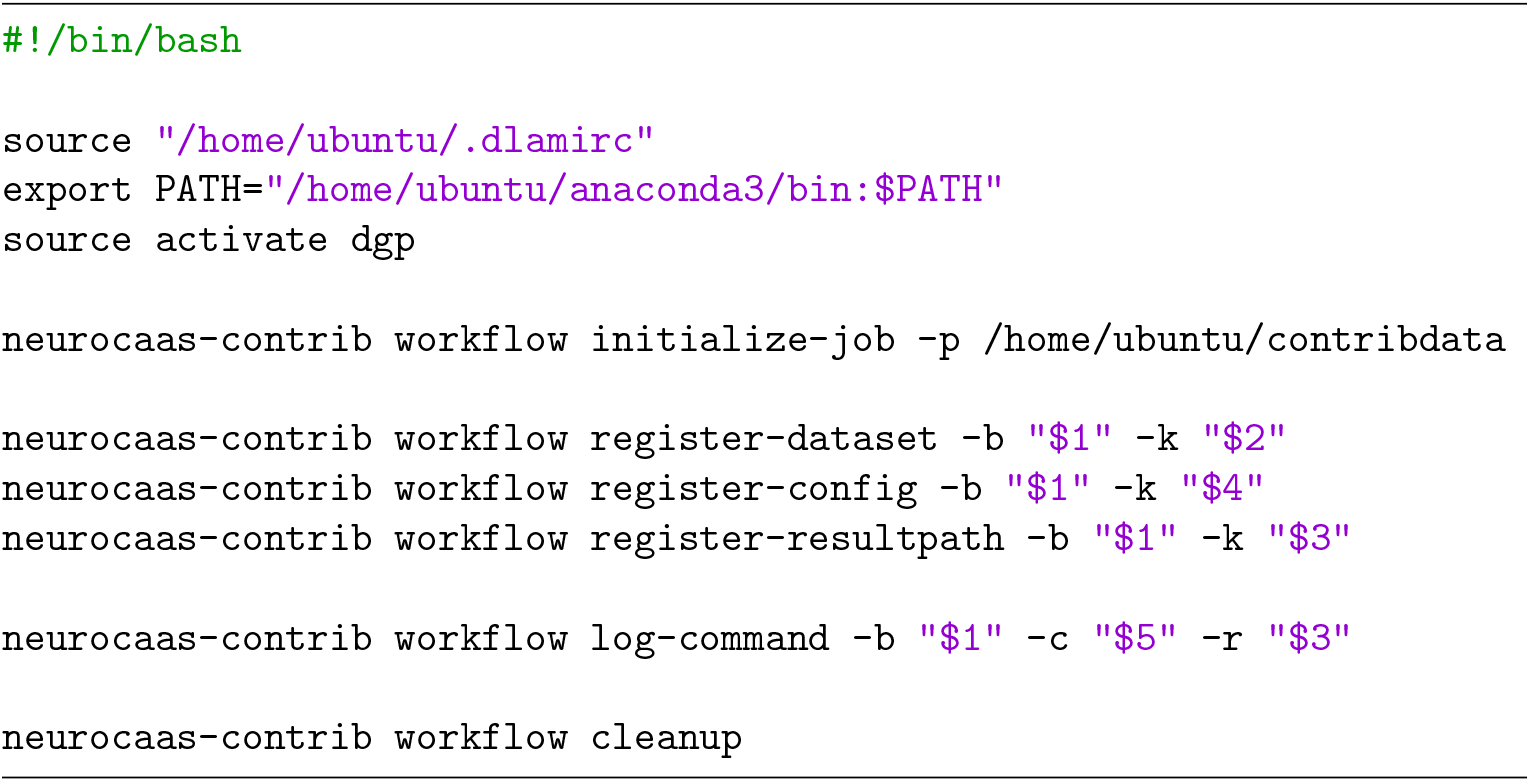
Main script called by NeuroCAAS to trigger workflow runs. The command neurocaas-contrib triggers developer tools. Declares variables referenced in Figure 12.

#### 5.1.1 Source Repo

The Platform section gives an overview of how NeuroCAAS encodes individual analyses into blueprints, and deploys them into full infrastructure stacks, following the principle of Infrastructure-as-Code (IaC). This section presents blueprints in more depth and show how the whole NeuroCAAS platform can be managed through IaC, encoding features such as user data storage, credentials, and logging infrastructure in code documents analogous to analysis blueprints as well. All of these code documents, together with code to deploy them, make up NeuroCAAS’s source repo. There is a one-to-one correspondence between NeuroCAAS’s source repo and infrastructure components: deploying the source repo provides total coverage of all the infrastructure needed to analyze data on NeuroCAAS (Figure 11, bottom).

Within the source repo, each NeuroCAAS blueprint (see Figure 14 for an example) is formatted as a JSON document with predefined fields. The expected values for most of these fields identify a particular cloud resource, such as the ID for an immutable analysis environment, or a hardware identifier to specify an instance within the resource bank (Lambda.LambdaConfig.AMI and Lambda.LambdaConfig.INSTANCE_TYPE in Figure 14, respectively). Upon deployment, these fields determine the creation of certain cloud resources: AWS EC2 Amazon Machine Images in the case of IAE IDs, and AWS EC2 Instances in the case of hardware identifiers. One notable exception is the protocol specifying behavior of a corresponding NeuroCAAS job manager (Lambda.CodeUri and Lambda.Handler in Figure 14). Instead of identifying a particular cloud resource, each blueprint’s protocol is a python module within the source repo that contains functions to execute tasks on the cloud in response to user input. The ability to specify protocols in python allows NeuroCAAS to support the complex workflows shown in Figure 8. Job managers are deployed from these protocols as AWS Lambda functions that execute the protocol code for a particular analysis whenever users submit data and parameters.

**Figure 14:**
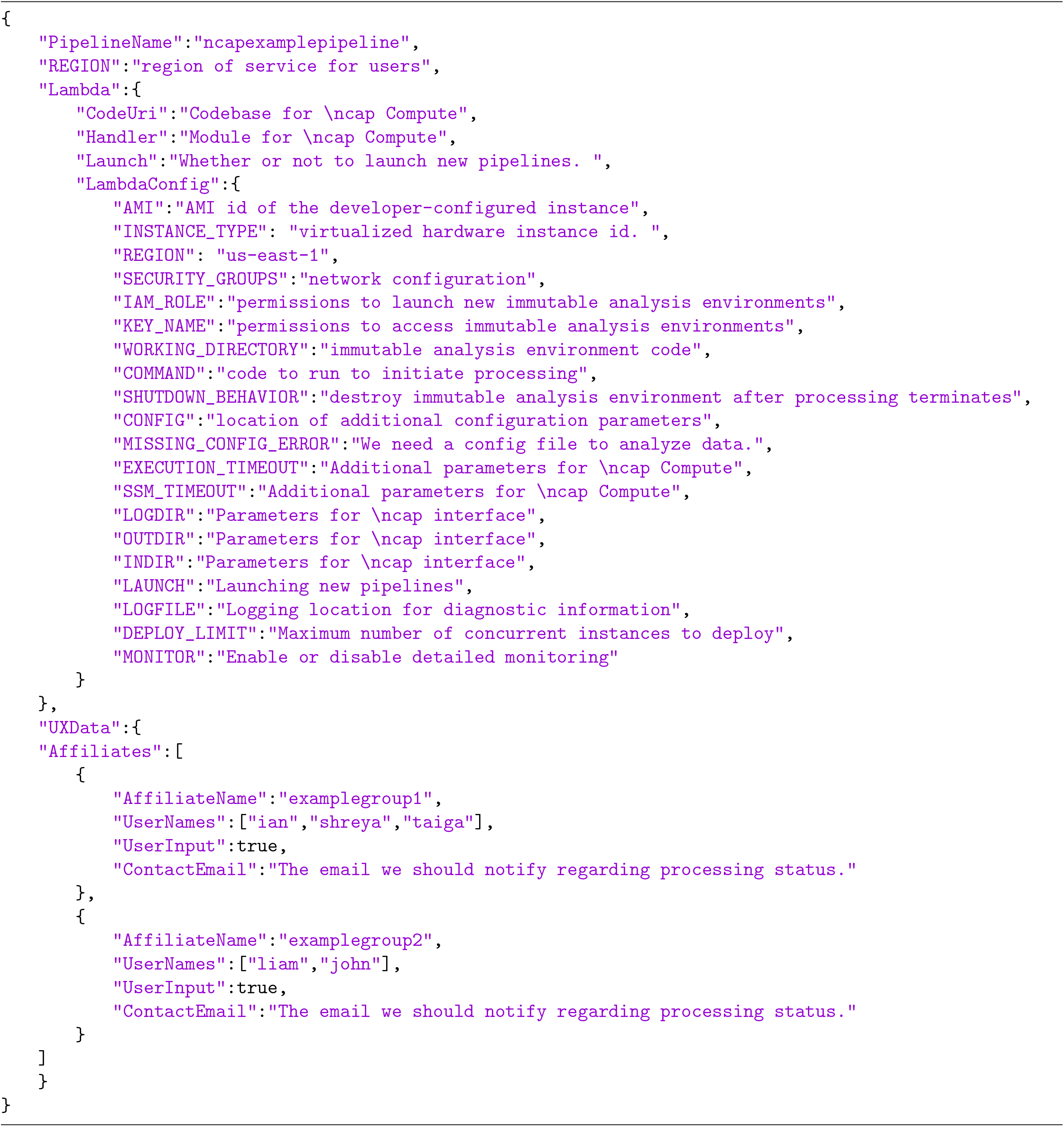
NeuroCAAS blueprint template declaring all relevant resources. Immutable Analysis Environments can be defined from Variables in the Lambda.LambdaConfig field, the job manager protocol is defined in Lambda.CodeUri and Lambda.Handler. Users and permissions are defined in UXData.

Another major aspect of NeuroCAAS’s source repo that is not discussed in the Platform section is the management of individual users. NeuroCAAS applies the same IaC principles to user creation and management as it does to individual analyses. To add a new user to the platform, NeuroCAAS first creates a corresponding user profile in the source repo (Figure 11, right), that specifies user budgets, creates private data storage space, generates their (encrypted) security credentials, and identifies other users who they collaborate with. Users resources are created using the AWS Identity and Accesss Management (IAM) service.

#### 5.1.2 Contrib and Interface Repos

Given only the NeuroCAAS source repo, analyses can be hosted on the NeuroCAAS platform and new users can be added to the platform simply by deploying the relevant code documents. However, interacting directly with resources provided by the NeuroCAAS source repo can be challenging for both analysis users and developers. For developers, the steps required to fill in a new analysis blueprint may not be clear, and the scripting steps necessary within an IAE to retrieve user data and parameters requires knowledge of specific resources on the Amazon Web Services cloud. For users, the NeuroCAAS source repo on its own does not support an intuitive interface or analysis documentation, requiring users to interact with NeuroCAAS through generic cloud storage browsers, forcing them to engage in tedious tasks like navigating file storage and downloading logs before examining them. Collectively, these tasks lower the accessibility that is a key part of NeuroCAAS’s intended design. To handle these challenges, we created two additional code repositories, the NeuroCAAS contrib repo and interface repo, for developers and users, respectively.

The NeuroCAAS contrib repo supports a command line tool and python code to streamline the process of developing and creating new NeuroCAAS analyses. During the development process, the NeuroCAAS contrib repo can create infrastructure stacks independently of input-triggered job managers for a limited time, allowing developers to build and test IAEs interactively on powerful hardware instances (Figure 11, bottom right), and populate the analysis blueprint as they go. Then, when a new analysis is ready to be used on NeuroCAAS, the NeuroCAAS contrib repo automatically versions the entire source repository after integrating and deploying the new blueprint, generating a unique analysis version ID. All NeuroCAAS analyses can be updated only by directly editing blueprints, and blueprints are assigned a new analysis version ID every time that they are updated. By enforcing a tight correspondence between blueprints and analyses, we ensured the reproducibility of all analyses conducted via NeuroCAAS, regardless of ongoing updates to the underlying infrastructure or algorithm (Figure 11, top right). With an analysis version ID, it is possible to replicate results that were generated with older versions of some analysis algorithm, making this a particularly useful feature for users processing data with an analysis that is still actively being developed. The NeuroCAAS contrib repo contains a detailed guide for developers to get started with NeuroCAAS.

The NeuroCAAS interface repo supports the website interface to NeuroCAAS, hosted at www.neurocaas.org. In addition to providing documentation and a simpler user interface, (Figure 11, bottom left) the interface repo automatically creates and deploys user profiles when users sign up, significantly increasing the potential scale of the platform (Figure 11, top left). This website based user credentialing system can be referenced by other user interfaces as well, as is done in https://github.com/jcouto/wfield. If users wish to share analysis access and data with other users, they can also use the website to create and request unique “group codes” at sign up, that they can use to invite other users into the same group. Doing so allows them to easily share analysis access with others.

### 5.2 Novel Analyses

For each novel analysis, we provide some details on its component infrastructure stacks, as well as details on relevant development outside the NeuroCAAS framework we have already presented.

#### 5.2.1 Widefield Imaging

The Widefield Calcium Imaging analysis that we present involves two independent infrastructure stacks, with the second taking as input the results of the first. The first infrastructure stack performs motion correction, denoising, compression, and hemodynamic correction, and is performed on an instance with 64 virtual cores (further infrastructure details are identical to the “PMD” row of Table 2). The second infrastructure stack performs demixing of denoised, corrected widefield imaging data, and is performed on an instance with a Tesla V100 GPU (further infrastructure details are identical to the “LocaNMF” row of Table 2). In addition to these two infrastructure stacks, we developed a custom graphical user interface (available for download at https://github.com/jcouto/wfield). This user interface integrates with the credentials generated for users on the NeuroCAAS website, allowing users who have signed up via the website to use the GUI with an existing account. The GUI hosts a number of initialization steps on the user’s local machine, involving selection of parameters and alignment of data to landmarks on a given brain atlas. The GUI is also able to upload data directly to NeuroCAAS cloud storage, submit jobs, and monitor their progress. Next, the GUI is able to detect when the first step of processing is completed, and submits the relevant results files as input to the second step, mimicking the steps a user would take manually to manage this process. Finally, when all processing is complete the GUI retrieves analysis results back to the user’s local machine. For more details on implementation of each analysis step, please see Couto et al. (2020).

**Table 2:** Infrastructure details for benchmarked algorithms. Job Monitor refers to the mechanisms used to track the status of ongoing jobs. Resource Usage refers to the hardware diagnostics tracked by NeuroCAAS. Version Control refers to the version control mechanisms used to maintain fidelity of core analysis code. Packages refers to the mechanisms used to handle analysis dependencies.

| NEUROCAAS Benchmarked Analyses |  |  |  |  |  |  |  |  |  |
| --- | --- | --- | --- | --- | --- | --- | --- | --- | --- |
| Analysis Name | Storage | Memory | GPU | CPU count | OS | Job Monitor | Resource Usage | Version Control | Packages |
| CaImAn | 500 GB SSD | 275 GB | N/A | 64 vCPU<sup>1</sup> | Ubuntu 16.04 (Linux HVM) | stdout | CPU Utilization | Github | pip requirements file |
| DeepLabCut | 200 GB SSD | 65 GB | Nvidia Tesla K80 | 4 vCPU<sup>2</sup> | Ubuntu 16.04 (Linux HVM) | stdout | CPU Utilization | Github | conda environment file |
| PMD | 75 GB SSD | 275 GB | N/A | 64 vCPU<sup>1</sup> | Ubuntu 18.04 (Linux HVM) | stdout | CPU Utilization | Github | Conda Package |
| LocaNMF | 75 GB SSD | 65 GB | Nvidia Tesla V100 | 8 vCPU<sup>3</sup> | Ubuntu 16.04 (Linux HVM) | stdout | CPU Utilization | Github | Conda Package |

<sup>1</sup> Intel Xeon Platinum 8000 series (Skylake-SP): https://aws.amazon.com/ec2/instance-types/m5/
<sup>2</sup> Intel Broadwell (AWS): https://aws.amazon.com/blogs/aws/new-p2-instance-type-for-amazon-ec2-up-to-16-gpus/
<sup>3</sup> Intel Xeon E5-2686v4: https://aws.amazon.com/blogs/aws/new-amazon-ec2-instances-with-up-to-8-nvidia-tesla-v100-gpus-p3/

#### 5.2.2 Ensemble Markerless Tracking

The deep ensembling analysis that we present is also performed is two separate infrastructure stacks, but both the initial training and the consensus output generation steps are performed on the same type of infrastructure. In both cases, we use an instance equipped with a Tesla V100 GPU, otherwise identical to the infrastructure shown in the DeepLabCut row of Table 2). We trained DeepGraphPose with the default training settings provided in the file run_dgp_demo.py, on the “twomice-top-down” data from the DeepGraphPose paper (Wu et al., 2020). That paper provides full videos of analysis of this dataset using a single DeepGraphPose model. To enable ensembling, we built a separate set of ensembling tools that work with DeepGraphPose (Wu et al., 2020) - they can be found at https://github.com/cunningham-lab/neurocaas_ensembles. In order to create a consensus output, we averaged the confidence maps from each model in an ensemble in the following way: Given a set of N trained DGP networks, *φ_i_, i* ∈ 1…*N*, and a video frame, 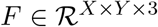. Assume that the network has been trained to track a single body part (the general case follows immediately), and take the scoremap outputs (unnormalized likelihoods) on this image from the output convolutional layer, denoted 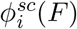, where each scoremap 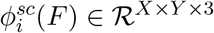. These scoremap outputs are unnormalized likelihoods representing the probability that the body part of interest is located in any individual pixel of the image. Then, we can compute the mean scoremap for a given image as:

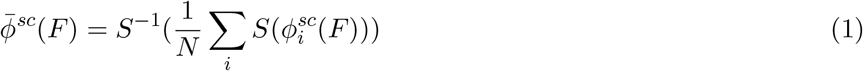

Where *S* is the elementwise sigmoid function. The consensus output is then calculated from the softargmax function of this mean scoremap.

Furthermore, to calculate the rmse error, we use the following metric: Assume we have detections for all of the test frames in a video as a tensor, 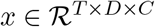, with entries *x_tdc_*, where t represents the frame index, d the part index, and c the coordinate ∈ [*x, y*]. Likewise, we have groundtruth data *g* with entries *g_tdc_* of the same dimension. Then the error is calculated as follows:

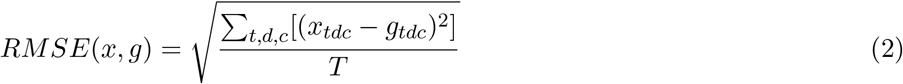

Details and implementation can be found in the repository https://github.com/cunningham-lab/neurocaas_ensembles, and the full analysis is available for use at http://neurocaas.org/analysis/14.

### 5.3 Benchmarking algorithms on NeuroCAAS

For each analysis currently on NeuroCAAS, the specific infrastructure choices in the corresponding blueprint (Figure 11, right) are given in Table 2. To meaningfully benchmark NeuroCAAS against current standards, we simulated corresponding *local* infrastructure. Local infrastructure was also built on AWS, and spans resources comparable to personal hardware and cluster compute, depending on the use case (see Table 3). As a general guideline, we chose local infrastructure representatives that would reasonably be available to a typical researcher, unless the datasets we considered required more powerful resources. To account for the diversity of resources available to neuroscience users, we offer alternative quantifications to those presented in Figure 10 in the supplementary methods (see Figure 15), and make performance quantification data and calculations available to users who would like to compare to their own infrastructure through a custom tool on our project repository (see README: https://github.com/cunningham-lab/neurocaas).

**Table 3:** Details of infrastructure used to simulate local processing. The column labels mirror those in Table 2

| NEUROCAAS Local Simulation |  |  |  |  |  |  |  |  |  |
| --- | --- | --- | --- | --- | --- | --- | --- | --- | --- |
| Analysis Name | Storage | Memory | GPU | CPU count | OS | Job Monitor | Resource Usage | Version Control | Packages |
| CaImAn | 500 GB SSD | 17 GB | N/A | 4 vCPU<sup>1</sup> | Ubuntu 16.04 (Linux HVM) | stdout | None | github | Pip requirements file |
| DeepLabCut | 200 GB SSD | 65 GB | Nvidia Tesla K80 | 4 vCPU<sup>2</sup> | Ubuntu 16.04 (Linux HVM) | stdout | None | github | Conda Package |
| PMD | 75 GB SSD | 131 GB | N/A | 16 vCPU<sup>3</sup> | Ubuntu 18.04 (Linux HVM) | stdout | None | Github | Conda Package |
| LocaNMF | 75 GB SSD | 65 GB | Nvidia Tesla K80 | 4 vCPU<sup>2</sup> | Ubuntu 16.04 (Linux HVM) | stdout | None | github | Conda Package |

<sup>1</sup> AMD EPYC 7000 Series: https://aws.amazon.com/ec2/instance-types/m5/
<sup>2</sup> Intel Broadwell (AWS): https://aws.amazon.com/blogs/aws/new-p2-instance-type-for-amazon-ec2-up-to-16-gpus/
<sup>3</sup> Intel Xeon E5 Broadwell Processors: https://aws.amazon.com/blogs/aws/new-next-generation-r4-memory-optimized-ec2-instances/

**Figure 15:**
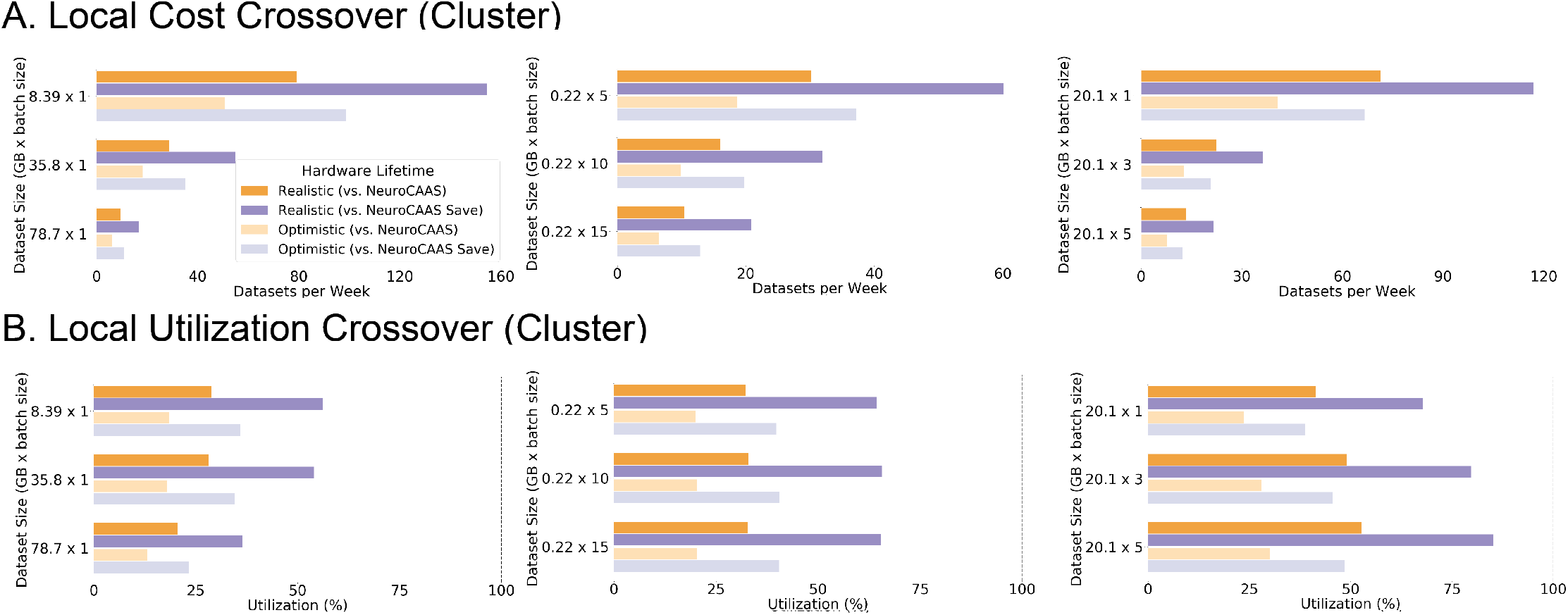
Alternative cost quantification of local infrastructure. A) provides Local Cost Crossover Crossover for these resources priced as cluster compute resources, priced according to Amazon AWS’s TCO calculator. B) provides the same for Local Utilization.

For each analysis that we benchmarked on NeuroCAAS, we chose three datasets of increasing size as representative use cases of the algorithms in question. The size differences of these datasets reflect the diversity of potential use cases among different users of the same algorithm. The CaImAn benchmarking data consists of datasets N.02.00, J_123, J_115 from the data shared with the CaImAn paper (Giovannucci et al., 2019). Benchmark analysis is based on a script provided to regenerate Figure 10 of the CaImAn paper. Note that although this data could be batched, we choose to maintain all three datasets as contiguous wholes. DeepLabCut benchmarking data consists of behavioral video capturing social interactions between two mice in their home cage. Data is provided courtesy of Robert C. Froemke and Ioana Carcea, as analyzed and presented in Carcea et al. (2019). Data processing consisted of analyzing these videos with a model that had previously been trained on other images from the same dataset. The same dataset was used to benchmark PMD and LocaNMF as a single analysis pipeline with two discrete parts. Input data consist of the dataset (“mSM30”), comprising widefield calcium imaging data videos, provided courtesy of Simon Musall and Anne Churchland, as used in Musall et al. (2019) and Saxena et al. (2020). The full dataset is available in a denoised format at http://repository.cshl.edu/id/eprint/38599/. Data processing on NeuroCAAS consisted of first processing the raw videos with PMD, then passing the resulting output to LocaNMF. Further details on the datasets used can be found in Table 5.

We split the time taken to run analyses on NeuroCAAS into two separate quantities. First, we quantified the time taken to upload data from local machines to NeuroCAAS, denoted as NeuroCAAS (Upload) in Figure 10. This time depends upon the specifics of the internet connection that is being used. It is also a one time cost: once data is uploaded to NeuroCAAS, it can be reanalyzed many times without incurring this cost again. Upload times were measured from the same NeuroCAAS interface made available to the user. (This upload time was skipped in the quantification of local processing time.) Second, we automatically quantified the total time elapsed between job submission and job termination, when results have been delivered back to the end user in the NeuroCAAS interface (denoted as NeuroCAAS (Compute) in Figure 10) via AWS native tools (see **Supplementary Methods** for details, and use of this data for Figure 6). Local timings were measured on automated portions of workflow in the same manner as NeuroCAAS (Compute).

We quantified the cost of running analysis on NeuroCAAS by enumerating costs of each of the AWS resources used in the course of a single analysis. Costs can be found in Table 4. We provide the raw quantification data and corresponding prices in Table 4. To further reduce costs, we also offer the option to utilize AWS Spot Instances (dedicated duration); these are functionally identical to standard compute instances, but are provisioned for a pre-determined amount of time with the benefit of significantly reduced prices. We provide the estimated cost of running analyses with both of these options in Figure 10, with quantifications labeled “NeuroCAAS Save” corresponding to analyses run with dedicated duration spot instances, and those labeled “NeuroCAAS Std” corresponding to those run with standard instances. For more on Spot Instance price quantification, see **Supplementary Methods**.

**Table 4:** Pricing details for implemented algorithms

| Pricing List |  |  |
| --- | --- | --- |
| Resource | Metrics | Rate |
| EC2 (Compute) | Time | Hardware Dependent, Fluctuates |
| Lambda (Workflow)<sup>1</sup> | Data Size × Time | $\$1.66667 \times 1e-5$ per GB-second |
| S3 (Data Transfer Out)<sup>2</sup> | Data Size | \$0.09 per GB |

<sup>1</sup> AWS Lambda is also priced for number of requests, but this is a negligible cost for a single analysis run.
<sup>2</sup> Data Transfer is only priced out of Amazon Web Services, i.e. in returning results to the end user.

**Table 5:** Details of the datasets used to benchmark performance. Sizes given for the three datasets tested for each pipeline shown. Dataset dimensionality labels are included in footnotes provided.

| Dataset Details |  |  |  |  |  |  |  |
| --- | --- | --- | --- | --- | --- | --- | --- |
| Analysis | Format | Size (Small) | Dim (Small) | Size (Medium) | Dim (Medium) | Size (Large) | Dim (Large) |
| CaImAn | zipped tiff | 8.39GB | $8000 \times 512 \times 512^1$ | 35.84GB | $41000 \times 458 \times 477^1$ | 78.70GB | $90000 \times 463 \times 472^2$ |
| DeepLabCut | mpeg | $5 \times 214.8\text{MB}$ | $5 \times 36000 \times 340 \times 420^3$ | $10 \times 214.8\text{MB}$ | $10 \times 36000 \times 340 \times 420^3$ | $15 \times 214.8\text{MB}$ | $15 \times 36000 \times 340 \times 420^3$ |
| PMD + LocaNMF | numpy array | $1 \times 20.1\text{GB}$ | $[500 \times 600 \times 1697, 1697 \times 8979]^4$ | $3 \times 20.1\text{GB}$ | $[500 \times 600 \times 1697, 1697 \times 8979];$<br>$[500 \times 600 \times 1652, 1652 \times 9000];$<br>$[500 \times 600 \times 2298, 2298 \times 8988]^4$ | $5 \times 20.1\text{GB}$ | $[500 \times 600 \times 1697, 1697 \times 8979];$<br>$[500 \times 600 \times 1652, 1652 \times 9000];$<br>$[500 \times 600 \times 2298, 2298 \times 8988];$<br>$[500 \times 600 \times 2304, 2304 \times 8992];$<br>$[500 \times 600 \times 1910, 1910 \times 8952]^4$ |

1 [Time × X × Y] at 7 hz Giovannucci et al. (2019)
2 [Time × X × Y] at 30 hz Giovannucci et al. (2019)
3 [Batch × Time × X × Y ] at 30 hz
4 [X × Y × Rank, Rank × Time] at 30 Hz

With simulated local infrastructures on AWS in hand, costs were calculated by pricing analogous computing resources as if the user had purchased them for a personal workstation, or as if they had been allocated to the user on an on-premises cluster (Table 8). In Figure 10, we assume that the local infrastructures considered are hosted on typical local laptop or desktop computing resources, supplemented with the resources necessary to run analyses as they were done on NeuroCAAS (additional storage, memory, GPU, etc), while maintaining approximate parity in processor power. We referred to (Morey and Nambiar, 2009) to convert pricetag costs of local machines to Equivalent Annual Costs, i.e. the effective cost per year if we assume our local machines will remain in service for a given number of years, as our implementation of a TCO calculation (as is often done in industry). Given a price tag cost *x_local_,* an assumed lifetime *n,* an annuity rate *r,* and *c_s_*(*n*) defined as the estimated annual cost of local machine support given a lifetime n, we follow Mahvi and Zarfaty (2009), Morey and Nambiar (2009) in calculating the Equivalent Annual Cost as:

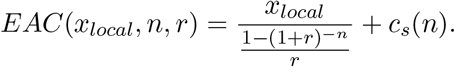

Here *c_s_*(*n*) is provided in the cited paper (Morey and Nambiar, 2009), estimated from representative data across many different industries. The denominator of the first term is an annuity factor. We consider two different values for n, which we label as “realistic” (2 years) and “optimistic” (4 years) in the text. In industry, 3-4 years is the generally accepted optimal lifespan for computers, after which support costs outweigh the value of maintaining an old machine (“Pilot Study”, 2004, Mahvi and Zarfaty, 2009, Morey and Nambiar, 2009). Some have argued that with more modern hardware, the optimal refresh cycle has shortened to 2 years (J.Gold Associates LLC, 2014). By providing quantifications assuming two and four year refresh cycle, we consider the short and long end of this generally discussed optimal range.

Given a per-dataset NeuroCAAS cost *x*_NeuroCAAS_, we further derive the Local Cost Crossover (LCC), the threshold weekly data analysis rate at which it becomes cost-effective to buy a local machine. The LCC is given by:

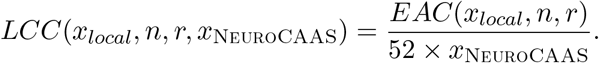

Furthermore, given the per-dataset local analysis time, we can estimate the corresponding Local Utilization Crossover (LUC). The LUC considers the LCC in the context of the maximal achievable data analysis rate on local infrastructure as calculated in the previous section. If the time taken to analyze a dataset on a local machine is given by *t_local_* (in seconds), The LUC is given by:

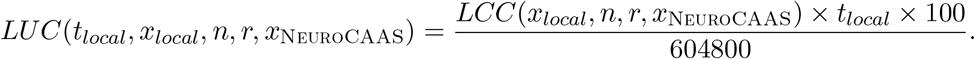

### 5.4 Survey of Analyses and Platforms

We characterized data analysis infrastructure as consisting of three hierarchical parts (Dependencies, System, Hardware), segmented consistently with infrastructure descriptions referenced elsewhere (Demchenko et al., 2013, Zhou et al., 2016). In several different subfields of neuroscience, we then selected 10 recent or prominent analysis techniques, and asked how they fulfilled each component of data analysis infrastructure to generate Figure 1D. We denoted a particular infrastructure component as supported if it is referenced in the relevant installation and usage guides as being provided in a reliable, automated manner (e.g., automatic package installation via pip). Survey details are provided in tables 6, 7. We addressed the question of how data analyses are installed and used with these surveys in the tradition of the open source usability literature. Surveys such as these are standard methodology in this field, which relies upon empirical data from studies of user’s usage habits (Nichols et al., 2001, Zhao and Deek, 2005), developer sentiment (Terry et al., 2010), and observation of user-developer interactions via platforms like Github (Cheng and Guo, 2018).

**Table 6:** Infrastructure support for Calcium Imaging Algorithms. Labels mirror those in Table 2.

| Calcium Imaging |  |  |  |  |  |  |  |  |  |  |
| --- | --- | --- | --- | --- | --- | --- | --- | --- | --- | --- |
| Algorithm Name | Publication | Software Version | Package Version | OS config | Batch Scripting | Other Processes | Storage | Memory | GPU | CPU |
| CaImAn | Giovannucci et al. 2017 | ✓ | ✓ | X | ✓ | X | X | X | X | X |
| CNMF-E | Zhou et al. 2018 | ✓ | ✓ | X | X | X | X | X | X | X |
| Suite2p | Pachitariu et al. 2017 | ✓ | ✓ | X | ✓ | X | X | X | X | X |
| ABLE | Reynolds et al. 2017 | ✓ | ✓ | X | ✓ | X | X | X | X | X |
| SCALPEL | Petersen et al. 2018 | ✓ | ✓ | X | X | X | X | X | X | X |
| Min1PIPE | Lu et al. 2018 | ✓ | ✓ | X | ✓ | X | X | X | X | X |
| SamuROI | Rueckl et al. 2017 | ✓ | X | X | X | X | X | X | X | X |
| Romano | Romano et al. 2017 | ✓ | ✓ | X | X | X | X | X | X | X |
| FISSA | Keemink et al. 2018 | ✓ | ✓ | X | ✓ | X | X | X | X | X |
| OASIS | Friedrich et al. 2017 | ✓ | ✓ | X | X | X | X | X | X | X |
| Percentage Supporting |  | 90% | 80% | 0% | 50% | 0% | 0% | 0% | 0% | 0% |

**Table 7:** Infrastructure support for Behavioral Quantification Algorithms. Labels mirror those in Table 2.

| Behavioral Quantification. |  |  |  |  |  |  |  |  |  |  |
| --- | --- | --- | --- | --- | --- | --- | --- | --- | --- | --- |
| Algorithm Name | Publication | Software Version | Package Version | OS config | Batch Scripting | Other Processes | Storage | Memory | GPU | CPU |
| DeepLabCut | Mathis et al. 2018 | ✓ | ✓ | ✓ | X | X | X | X | X | X |
| DeepFly3D | Günel et al. 2019 | ✓ | ✓ | X | X | X | X | X | X | X |
| JAABA | Kabra et al. 2012 | ✓ | ✓ | X | X | X | X | X | X | X |
| Ctrax | Branson et al. 2009 | ✓ | ✓ | X | ✓ | X | X | X | X | X |
| DeepPoseKit | Graving et al. 2019 | ✓ | X | X | X | X | X | X | X | X |
| Ethovision | — | ✓ | ✓ | X | X | X | X | X | X | X |
| APT | — | ✓ | ✓ | ✓ | X | X | X | X | X | X |
| bonsai | Lopes et al. 2015 | X | ✓ | X | X | X | X | X | X | X |
| Miceprofiler | de Chaumont et al. 2012 | ✓ | ✓ | X | X | X | X | X | X | X |
| LEAP | Pereira et al. 2018 | ✓ | ✓ | X | X | X | X | X | X | X |
| Percentage Supporting |  | 90% | 90% | 20% | 10% | 0% | 0% | 0% | 0% | 0% |

**Table 8:** Instance and hardware cost details for local cost comparisons. Estimated Price tag prices as of May 3rd, 2020. Price tag estimation of workstation style hardware was based on market prices chosen to reflect the infrastructure implementation as given in Table 3, in particular, CPU make. Estimation of cluster style hardware cost was based on the AWS TCO calculator (https://awstcocalculator.com), as of January 25th, 2020, incorporating the total server hardware cost (undiscounted) and acquisition cost of SAN storage.

| Cost (Local) |  |  |  |  |  |  |
| --- | --- | --- | --- | --- | --- | --- |
| Algorithm Name | vCPU count | GPU | Memory | Storage Capacity | Workstation Price, US Dollars (Estimated Price Tag Cost) | Cluster Price, US Dollars (Estimated Price Tag Cost from Amazon TCO Calculator) |
| CaImAn | 4 | N/A | 16 GiB | 500 GB | 1618<sup>2</sup> | 1499+1000 |
| DeepLabCut | 4 | Tesla K80 | 61GiB | 200 GB | 3120<sup>3</sup> | 1701+400+1555<sup>5</sup> |
| PMD + LocaNMF <sup>1</sup> | 16 | Tesla K80 | 122 GiB | 150 GB | 5436<sup>4</sup> | 10836+300+1555<sup>5</sup> |

<sup>1</sup> Cost for PMD and LocaNMF refers to hardware cost for a local instance that can account for processing done on both.
<sup>2</sup> https://www.newegg.com/p/1TS-000D-052P6
<sup>3</sup> https://www.newegg.com/p/1VK-001E-1SVY3?Item=9SIADB38AG7178&Description=1080%20ti%20workstation%2064%20gB%204%20core&cm_re=1080_ti_workstation_64_gB_4_core_-_1VK-001E-1SVY3_-_Product
<sup>4</sup> https://www.newegg.com/p/1VK-001E-1A6V1
<sup>5</sup> https://www.vgastore.com/2023019/hp-j0g95a-tesla-k80-24gb-384-bit-gddr5-pci-e-3-0-x16-graphics-card

To generate Figure 7, we first quantified the traffic and infrastructure experienced by individual analyses by examining their Github pages, and taking the maximum of the number of forks, stars, and watchers, or downloads if listed as well as the listed hardware requirements of each analysis (numbers as of September 2020). We then overlaid several exemplar platforms based on the analyses that they supported, as well as restrictions based on the accessibility and scale requirements imposed by each (local hardware, limitation to one analysis at a time), taking care to include analyses that the platforms supported in practice.

## 6 Acknowledgements

We thank Ioana Carcea and Robert C. Froemke for the use of benchmarking data for DeepLabCut, Simon Musall and Anne Churchland for the use of benchmarking data for Penalized Matrix Decomposition and LocaNMF, Erica Rodriguez and C. Daniel Salzman for the use of data to demonstrate ensemble markerless tracking, Jian Wang and Yakov Stern for helpful discussions on NeuroCAAS’s architecture and extensions, Peter Lee, Jackson Loper and Shuonan Chen for development of additional algorithms, Selmaan Chettih for suggesting ensembling of pose tracking, Zahra Adahman, Ioana Carcea, Claire Everett, Andres Bendesky, Andres Villegas, Franck Polleux, Vivek Athalye, Darcy Peterka, and Avner Wallach for discussion and feedback on the use of NeuroCAAS during development, and Dan Biderman and Danil Tyulmankov for useful references and discussions of benchmarking. T.A. is supported by NIH training grant 2T32NS064929-11. S.S. is supported by the Swiss National Science Foundation P2SKP2_178197 and 5U19NS104649. E.K.B. is supported by NIH training grant 2T32NS064929-11, NIH U19NS107613, NSF DBI-1707398, and the Gatsby Charitable Foundation (Gatsby Charitable Foundation GAT3708). J.P.C. is supported by Simons 542963. L.P. is funded by IARPA MICRONS D16PC00003, NIH 5U01NS103489, 5U19NS104649, 5U19NS107613, 1UF1NS107696, 1UF1NS108213, 1RF1MH120680, DARPA NESD N66001-17-C-4002 and Simons Foundation 543023. L.P. and J.P.C. are supported by NSF Neuronex Award DBI-1707398.

## 7 Author Contributions

T.A. and J.P.C. conceived of the project. T.A. designed the infrastructure platform with input from all authors. S.S., I.K., S.L.K, R.G. J.Z and T.A. developed analyses to work on the platform (with supervision from L.P. and J.P.C.), and S.S., I.K., and T.A. collected the data shown in Figure 4. J.C., S.S., I.K. developed code for WFCI analysis and GUI, E.K.B. and T.A. developed code for ensembled markerless tracking with supervision from L.P., and J.B. developed and maintained the website. T.A., L.P. and J.P.C. wrote the paper with input from all authors.

## 9 Supplementary Methods

### 9.1 Managing users from user profiles

On NeuroCAAS, users resources were defined in code via JSON documents we call user profiles. New users were registered by filling in a corresponding user profile, which was then deployed to automatically generate storage space, dedicated login credentials, and permissions to use analyses for the user. The user profile is similar in format to the UXData segment of the blueprint as given in Figure 14, and can be found in the NeuroCAAS codebase online.

Deploying user profiles created a secure, virtualized storage location where users could store their data on NeuroCAAS before and in between analyses. Data storage on NeuroCAAS is shared within a user group (i.e. a lab), but private to all other parties. In Figure 10, NeuroCAAS Upload time refers to the time required to upload data from local machines to this storage- once uploaded, data can be deleted post analysis or maintained over the course of several analyses. Maintaining data post analysis cuts out NeuroCAAS Upload time on subsequent upload events.

User credentials are automatically generated upon new user sign up. Permissions to use analyses can be managed in a variety of ways. During the active development phase, permissions can be given to individual users by updating NeuroCAAS blueprints with the information of these newly added users, and redeploying the analysis in question- these analysis-centric permissions are best suited to analyses where one wishes to invite a small number of test users. Upon redeployment, the corresponding job manager begins monitoring this new user for analysis requests. Once analyses are stable, they can instead be accessed with user-centric permissions: when new user groups are created, they can have a pre-determined list of analyses that they will have access to from the outset. Importantly, both user centric and developer centric permissions schemes do not disrupt ongoing analysis jobs.

### 9.2 Automatic compute benchmarking

The duration of NeuroCAAS Compute and Local analysis time was recorded automatically with cloud native resource monitoring tools (AWS Lambda, AWS Cloudwatch Events, and AWS S3). These tools automatically recorded the creation and destruction of instances, and recorded the relevant timestamps at millisecond resolution. These monitoring tools were also managed via NeuroCAAS blueprints, and their design can be found in the same blueprint codebase. The same tools were used to calculate the usage data shwon in Figure 6. We do not disclose user data at an individual level, but developers can generate the same figures for users of their own analysis by using the NeuroCAAS contrib repo (specific instructions are given in the source repo’s README file).

### 9.3 Spot instance pricing

The virtualized hardware underlying a hardware instance can be provisioned at several different prices. We used AWS EC2 Spot Instance pricing to reduce costs, having known beforehands how long the analyses would take. At the moment, we depict prices based on spot instance availability in September 2019. Empirically, we observe that spot instance price fluctuations give standard deviations on the order of cents over a period of months (see source repo for experiments).

### 9.4 Analysis reproducibility

Because we designed analysis blueprint to be git versioned, we can reproduce the infrastructure and software configuration used to generate any analysis, up to the reliability of Amazon AWS. Since we returned identifying information about the blueprint to the user in a certificate along with configuration parameters, data is the only portion of an analysis that must be maintained to ensure perfect analysis reproducibility. We update blueprints based on pull requests issued through the source repo’s Github page, providing a centralized way to manage the state of the NeuroCAAS platform. Although not implemented here, AWS offers cheap, glacial storage that can be used to preserve data for long amounts of time under conditions of infrequent access, offering a feasible solution for guaranteed total analysis reproducibility on NeuroCAAS.

### 9.5 Alternative local crossovers

Because the instances offered on AWS are not wholly analogous to either personal hardware or cluster resources, we offer additional comparisons that span the range of prices.

Cluster pricing was calculated with the AWS TCO calculator https://awstcocalculator.com/#. We calculated the cost of infrastructure as a subset of the TCO provided by AWS. In particular, we calculated *x_local_* as the total server hardware cost (undiscounted) and acquisition cost of NAS storage, and the cost of a GPU, with additional yearly recurring costs *c_s_*(*n*) given by storage administration cost, server hardware maintenance cost, and IT Labor costs. We then calculated the LCC and LUC from these quantities as described in the Platform section.

The results of these quantifications are given in Figure 15.

### 9.6 AWS details

We provide further details on the AWS implementation of analyses used to generate time and cost data in Table 9.

**Table 9:** Instance and Amazon Machine Image (AMI) details for cost comparison of some implemented algorithms.

| NEUROCAAS AWS Specifics |  |  |  |
| --- | --- | --- | --- |
| Analysis Name | Instance (NEUROCAAS) | Instance (local) | AMI ID |
| CaImAn | m5.16xlarge | m5a.xlarge | ami-01dc867df8c05aa5a) |
| DeepLabCut | p2.xlarge | p2.xlarge | ami-00b1babeb8637f5c3) |
| PMD | m5.16xlarge | r4.4xlarge | ami-0007adf33fbcf0c1c) |
| LocaNMF | p3.2xlarge | p2.xlarge | ami-04ebe747c2e33038c) |

## References

Martin Abadi, Paul Barham, Jianmin Chen, Zhifeng Chen, Andy Davis, Jeffrey Dean, Matthieu Devin, Sanjay Ghemawat, Geoffrey Irving, Michael Isard, et al. Tensorflow: A system for large-scale machine learning. In 12th Symposium on Operating Systems Design and Implementation (OSDI16), pages 265–283, 2016.

Ademar Aguiar, Jessica Díaz, Rubén Almaraz, Jennifer Pérez, and Juan Garbajosa. DevOps in Practice – An Exploratory Case Study. In Proceedings of the 19th International Conference on Agile Software Development: Companion, XP ‘18, pages 1–3, 2018. ISBN 9781450364225. doi: 10.1145/3234152.3234199.

Robert A Amezquita, Aaron TL Lun, Etienne Becht, Vince J Carey, Lindsay N Carpp, Ludwig Geistlinger, Federico Martini, Kevin Rue-Albrecht, Davide Risso, Charlotte Soneson, et al. Orchestrating single-cell analysis with Bioconductor. Nature methods, pages 1–9, 2019.

Peter Amstutz, Michael R. Crusoe, Neboj sa Tijanic, Brad Chapman, John Chilton, Michael Heuer, Andrey Kartashov, John Kern, Dan Leehr, Herve Menager, Maya Nedeljkovich, Matt Scales, Stian Soiland-Reyes, and Luka Stojanovic. Common Workflow Language, v1.0, 2016. URL https://doi.org/10.6084/m9.figshare.3115156.v2.

Eleanor Batty, Josh Merel, Nora Brackbill, Alexander Heitman, Alexander Sher, Alan Litke, EJ Chichilnisky, and Liam Paninski. Multilayer recurrent network models of primate retinal ganglion cell responses. International Conference on Learning Representations, 2016.

Eleanor Batty, Matthew Whiteway, Shreya Saxena, Dan Biderman, Taiga Abe, Simon Musall, Winthrop Gillis, Jeffrey Markowitz, Anne Churchland, John P Cunningham, et al. BehaveNet: nonlinear embedding and Bayesian neural decoding of behavioral videos. In Advances in Neural Information Processing Systems, pages 15680–15691, 2019.

Sean R. Bittner, Agostina Palmigiano, Alex T. Piet, Chunyu A. Duan, Carlos D. Brody, Kenneth D. Miller, and John P. Cunningham. Interrogating theoretical models of neural computation with deep inference. bioRxiv, page 837567, 2019. doi: 10.1101/837567.

Joshua Bloch. Effective java (the java series). Prentice Hall PTR, 2008.

Alessio Paolo Buccino, Cole Lincoln Hurwitz, Samuel Garcia, Jeremy Magland, Joshua H Siegle, Roger Hurwitz, and Matthias H Hennig. SpikeInterface, a unified framework for spike sorting. eLife, 9:e61834, 2020. doi: 10.7554/elife.61834.

E Kelly Buchanan, Ian Kinsella, Ding Zhou, Rong Zhu, Pengcheng Zhou, Felipe Gerhard, John Ferrante, Ying Ma, Sharon Kim, Mohammed Shaik, et al. Penalized matrix decomposition for denoising, compression, and improved demixing of functional imaging data. arXiv preprint arXiv:1807.06203, 2018.

Jonathan B Buckheit and David L Donoho. Wavelab and reproducible research. In Wavelets and statistics, pages 55–81. Springer, 1995.

Business Intelligence. Pilot Study: Optimum Refresh Cycle and Method for Desktop Outsourcing. Technical report, Intel Business Center, 2004.

Ioana Carcea, Naomi Lopez Caraballo, Bianca J Marlin, Rumi Ooyama, Joyce M Mendoza Navarro, Maya Open-dak, Veronica E Diaz, Luisa Schuster, Maria I Alvarado Torres, Harper Lethin, et al. Oxytocin Neurons Enable Social Transmission of Maternal Behavior. bioRxiv, page 845495, 2019.

Anne E Carpenter, Thouis R Jones, Michael R Lamprecht, Colin Clarke, In Han Kang, Ola Friman, David A Guertin, Joo Han Chang, Robert A Lindquist, Jason Moffat, et al. CellProfiler: image analysis software for identifying and quantifying cell phenotypes. Genome biology, 7(10):R100, 2006.

Jeffrey C Carver, Sandra Gesing, Daniel S Katz, Karthik Ram, and Nicholas Weber. Conceptualization of a us research software sustainability institute (URSSI). Computing in Science \& Engineering, 20(3):4–9, 2018.

Chan Zuckerberg Initiative. Essential Open Source Software for Science (EOSS) - Chan Zuckerberg Initiative, 05 2019. URL https://chanzuckerberg.com/eoss/.

Fabrice De Chaumont, Stephane Dallongeville, Nicolas Chenouard, Nicolas Herve, Sorin Pop, Thomas Provoost, Vannary Meas-Yedid, Praveen Pankajakshan, Timothee Lecomte, Yoann Le Montagner, et al. Icy: an open bioimage informatics platform for extended reproducible research. Nature methods, 9(7):690, 2012.

Shuonan Chen, Jackson Loper, Xiaoyin Chen, Tony Zador, and Liam Paninski. BARcode DEmixing through Non-negative Spatial Regression (BarDensr). bioRxiv, page 2020.08.17.253666, 2020. doi: 10.1101/2020.08.17.253666.

Xiaoli Chen, Sunje Dallmeier-Tiessen, Robin Dasler, Sebastian Feger, Pamfilos Fokianos, Jose Benito Gonzalez, Harri Hirvonsalo, Dinos Kousidis, Artemis Lavasa, Salvatore Mele, et al. Open is not enough. Nature Physics, 15(2):113–119, 2019.

Jinghui Cheng and Jin LC Guo. How do the open source communities address usability and ux issues?: An exploratory study. In Extended Abstracts of the 2018 CHI Conference on Human Factors in Computing Systems,page LBW523, 2018.

Fernando Chirigati, Remi Rampin, Dennis Shasha, and Juliana Freire. ReproZip: Computational Reproducibility with Ease. In SIGMOD ‘16: Proceedings of the 2016 International Conference on Management of Data, 2016.

Joao Couto, Simon Musall, Xiaonan R Sun, Anup Khanal, Steven Gluf, Shreya Saxena, Ian Kinsella, Taiga Abe, John P Cunningham, Liam Paninski, and Anne K Churchland. Chronic, cortex-wide imaging of specific cell populations during behavior. arXiv, 2020.

Sharon M Crook, Andrew P Davison, and Hans E Plesser. Learning from the past: approaches for reproducibility in computational neuroscience. In 20 Years of Computational Neuroscience, pages 73–102. Springer, 2013.

Dandi Team. Dandi Archive, 2019. URL https://www.dandiarchive.org/.

Yuri Demchenko, Paola Grosso, Cees De Laat, and Peter Membrey. Addressing big data issues in scientific data infrastructure. In 2013 International Conference on Collaboration Technologies and Systems (CTS), pages 48–55, 2013.

Thomas G. Dietterich. Multiple Classifier Systems, First International Workshop, MCS 2000 Cagliari, Italy, June 21-23, 2000 Proceedings. Lecture Notes in Computer Science, pages 1–15, 2000. ISSN 0302-9743. doi: 10.1007/3-540-45014-9\_1.

David L Donoho. An invitation to reproducible computational research. Biostatistics, 11(3):385–388, 2010.

Editorial. Code share: Nature News & Comment, 10 2014. URL https://www.nature.com/news/code-share-1.16232.

Flywheel Exchange. Flywheel • Informatics Platform for Biomedical Research & Collaboration, 2019. URL https://flywheel.io/.

Stanislav Fort, Huiyi Hu, and Balaji Lakshminarayanan. Deep Ensembles: A Loss Landscape Perspective. arXiv, 2019.

Jeremy Freeman. Open source tools for large-scale neuroscience. Current opinion in neurobiology, 32:156–163, 2015.

Yuanjun Gao, Evan W Archer, Liam Paninski, and John P Cunningham. Linear dynamical neural population models through nonlinear embeddings. In Advances in neural information processing systems, pages 163–171, 2016.

Andrea Giovannucci, Johannes Friedrich, Matt Kaufman, Anne Churchland, Dmitri Chklovskii, Liam Paninski, and Eftychios A Pnevmatikakis. Onacid: Online analysis of calcium imaging data in real time. In Advances in neural information processing systems, pages 2381–2391, 2017.

Andrea Giovannucci, Johannes Friedrich, Pat Gunn, Jeremie Kalfon, Brandon L Brown, Sue Ann Koay, Jiannis Taxidis, Farzaneh Najafi, Jeffrey L Gauthier, Pengcheng Zhou, et al. CaImAn an open source tool for scalable calcium imaging data analysis. Elife, 8:e38173, 2019.

Tristan Glatard, Lindsay B. Lewis, Rafael Ferreira da Silva, Reza Adalat, Natacha Beck, Claude Lepage, Pierre Rioux, Marc-Etienne Rousseau, Tarek Sherif, Ewa Deelman, Najmeh Khalili-Mahani, and Alan C. Evans. Reproducibility of neuroimaging analyses across operating systems. Frontiers in Neuroinformatics, 9:12, 2015. doi: 10.3389/fninf.2015.00012.

Dan F. M. Goodman and Romain Brette. The Brian Simulator. Frontiers in Neuroscience, 3(2):192–197, 2009. ISSN 1662-4548. doi: 10.3389/neuro.01.026.2009.

Google Research. Colaboratory FAQ, 2017. URL https://research.google.com/colaboratory/faq.html#resource-limits.

Krzysztof Gorgolewski, Christopher D Burns, Cindee Madison, Dav Clark, Yaroslav O Halchenko, Michael L Waskom, and Satrajit S Ghosh. Nipype: a flexible, lightweight and extensible neuroimaging data processing framework in python. Frontiers in neuroinformatics, 5:13, 2011.

Krzysztof J Gorgolewski, Fidel Alfaro-Almagro, Tibor Auer, Pierre Bellec, Mihai Capota, M Mallar Chakravarty, Nathan W Churchill, Alexander Li Cohen, R Cameron Craddock, Gabriel A Devenyi, et al. BIDS apps: Improving ease of use, accessibility, and reproducibility of neuroimaging data analysis methods. PLoS computational biology, 13(3):e1005209, 2017.

Jacob M Graving, Daniel Chae, Hemal Naik, Liang Li, Benjamin Koger, Blair R Costelloe, and Iain D Couzin. DeepPoseKit, a software toolkit for fast and robust animal pose estimation using deep learning. eLife, 8:e47994, 2019. doi: 10.7554/elife.47994.

Klaus Greff, Aaron Klein, Martin Chovanec, Frank Hutter, and Jurgen Schmidhuber. The Sacred Infrastructure for Computational Research. In Proceedings of the 16th Python in Science Conference, 2017.

Yaroslav O. Halchenko and Michael Hanke. Open is Not Enough. Let’s Take the Next Step: An Integrated, Community-Driven Computing Platform for Neuroscience. Frontiers in Neuroinformatics, 6:22, 2012. doi: 10.3389/fninf.2012.00022.

Brooks Hanson, Andrew Sugden, and Bruce Alberts. Making data maximally available. Science, 331(6018):649, 2011.

Konrad Hinsen. Technical Debt in Computational Science. Computing in Science & Engineering, 17(6):103–107, 2015. ISSN 1521-9615. doi: 10.1109/mcse.2015.113.

Christina Hoffa, Gaurang Mehta, Tim Freeman, Ewa Deelman, Kate Keahey, Bruce Berriman, and John Good. On the use of cloud computing for scientific workflows. In 2008 IEEE fourth international conference on eScience, pages 640–645, 2008.

Michal Januszewski, Jörgen Kornfeld, Peter H. Li, Art Pope, Tim Blakely, Larry Lindsey, Jeremy Maitin-Shepard, Mike Tyka, Winfried Denk, and Viren Jain. High-precision automated reconstruction of neurons with flood-filling networks. Nature Methods, 15(8):605–610, 2018. ISSN 1548-7091. doi: 10.1038/s41592-018-0049-4.

Yaser Jararweh, Mahmoud Al-Ayyoub, Elhadj Benkhelifa, Mladen Vouk, Andy Rindos, et al. Software defined cloud: Survey, system and evaluation. Future Generation Computer Systems, 58:56–74, 2016.

J.Gold Associates LLC. Replacing Enterprise PCs: The Fallacy of the 3-4 Year Upgrade Cycle [White Paper]. Technical report, J.Gold Associates LLC, 2014.

Gary A Kane, Gonçalo Lopes, Jonny L Sanders, Alexander Mathis, and Mackenzie Mathis. Real-time, low-latency closed-loop feedback using markerless posture tracking. eLife, 9:e61909, 2020. doi: 10.7554/elife.61909.

Johannes Koster and Sven Rahmann. Snakemake—a scalable bioinformatics workflow engine. Bioinformatics, 28 (19):2520–2522, 2012.

Matthew Krafczyk, August Shi, Adhithya Bhaskar, Darko Marinov, and Victoria Stodden. Scientific Tests and Continuous Integration Strategies to Enhance Reproducibility in the Scientific Software Context. In Proceedings of the 2nd International Workshop on Practical Reproducible Evaluation of Computer Systems, pages 23–28, 2019.

Balaji Lakshminarayanan, Alexander Pritzel, and Charles Blundell. Simple and Scalable Predictive Uncertainty Estimation using Deep Ensembles. arXiv, 2016.

Esther Landhuis. Neuroscience: Big brain, big data. Nature, 541(7638):559–561, 2017. ISSN 0028-0836. doi: 10.1038/541559a.

Jin Hyung Lee, David E Carlson, Hooshmand Shokri Razaghi, Weichi Yao, Georges A Goetz, Espen Hagen, Eleanor Batty, EJ Chichilnisky, Gaute T Einevoll, and Liam Paninski. Yass: Yet another spike sorter. In Advances in neural information processing systems, pages 4002–4012, 2017.

Goncalo Lopes, Niccolo Bonacchi, Joao Frazao, Joana P Neto, Bassam V Atallah, Sofia Soares, Luis Moreira, Sara Matias, Pavel M Itskov, Patricia A Correia, et al. Bonsai: an event-based framework for processing and controlling data streams. Frontiers in neuroinformatics, 9:7, 2015.

Jeremy Magland, James J Jun, Elizabeth Lovero, Alexander J Morley, Cole Lincoln Hurwitz, Alessio Paolo Buccino, Samuel Garcia, and Alex H Barnett. SpikeForest, reproducible web-facing ground-truth validation of automated neural spike sorters. eLife, 9:e55167, 2020. doi: 10.7554/elife.55167.

John Mahvi and Avi Zarfaty. Using TCO to Determine PC Upgrade Cycles. Corporation, Intel, 2009.

Alexander Mathis, Pranav Mamidanna, Kevin M Cury, Taiga Abe, Venkatesh N Murthy, Mackenzie Weygandt Mathis, and Matthias Bethge. DeepLabCut: markerless pose estimation of user-defined body parts with deep learning. Nature Neuroscience, 21(9), 2018.

Zeeya Merali. Computational science: … Error. Nature, 467(7317):775–777, 2010. ISSN 0028-0836. doi: 10.1038/467775a.

Dirk Merkel. Docker: lightweight linux containers for consistent development and deployment. Linux journal, 2014(239):2, 2014.

Greg Miller. A Scientist’s Nightmare: Software Problem Leads to Five Retractions. Science, 314(5807):1856–1857, 2006. ISSN 0036-8075. doi: 10.1126/science.314.5807.1856.

Thomas P. Minka. From Hidden Markov Models to Linear Dynamical Systems, 1999.

Hatef Monajemi, Riccardo Murri, Eric Jonas, Percy Liang, Victoria Stodden, and David L Donoho. Ambitious Data Science Can Be Painless. arXiv preprint arXiv:1901.08705, 2019.

Timothy Morey and Roopa Nambiar. Using Total Cost of Owner-ship to Determine Optimal PC Refresh Lifecycles [White Paper]. Technical report, Wipro Ltd., 2009.

Kief Morris. Infrastructure as code: managing servers in the cloud.“ O’Reilly Media, Inc.”, 2016.

Simon Musall, Matthew T Kaufman, Ashley L Juavinett, Steven Gluf, and Anne K Churchland. Single-trial neural dynamics are dominated by richly varied movements. Nature neuroscience, 22(10):1677–1686, 2019.

David M Nichols, Kirsten Thomson, and Stuart Andrew Yeates. Usability and open-source software development. In CHINZ’01, pages 49–54, 2001.

Simon RO Nilsson, Nastacia L. Goodwin, Jia Jie Choong, Sophia Hwang, Hayden R Wright, Zane C Norville, Xiaoyu Tong, Dayu Lin, Brandon S. Bentzley, Neir Eshel, Ryan J McLaughlin, and Sam A. Golden. Simple Behavioral Analysis (SimBA) – an open source toolkit for computer classification of complex social behaviors in experimental animals. bioRxiv, page 2020.04.19.049452, 2020. doi: 10.1101/2020.04.19.049452.

Anna Nowogrodzki. How to support open source software and stay sane. Nature, 571:133–134, 2019.

Yaniv Ovadia, Emily Fertig, Jie Ren, Zachary Nado, D Sculley, Sebastian Nowozin, Joshua V Dillon, Balaji Lakshminarayanan, and Jasper Snoek. Can You Trust Your Model’s Uncertainty? Evaluating Predictive Uncertainty Under Dataset Shift. arXiv, 2019.

Marius Pachitariu, Nicholas A Steinmetz, Shabnam N Kadir, Matteo Carandini, and Kenneth D Harris. Fast and accurate spike sorting of high-channel count probes with KiloSort. In Advances in Neural Information Processing Systems, pages 4448–4456, 2016.

Marius Pachitariu, Carsen Stringer, Mario Dipoppa, Sylvia Schroder, L Federico Rossi, Henry Dalgleish, Matteo Carandini, and Kenneth D Harris. Suite2p: beyond 10,000 neurons with standard two-photon microscopy. Bioarxiv, page 061507, 2017.

Chethan Pandarinath, Daniel J O’Shea, Jasmine Collins, Rafal Jozefowicz, Sergey D Stavisky, Jonathan C Kao, Eric M Trautmann, Matthew T Kaufman, Stephen I Ryu, Leigh R Hochberg, et al. Inferring single-trial neural population dynamics using sequential auto-encoders. Nature methods, 15(10):805–815, 2018.

Liam Paninski and John Cunningham. Neural data science: accelerating the experiment-analysis-theory cycle in large-scale neuroscience. Current opinion in neurobiology, 50:232–241, 2018.

Nikhil Parthasarathy, Eleanor Batty, William Falcon, Thomas Rutten, Mohit Rajpal, EJ Chichilnisky, and Liam Paninski. Neural networks for efficient bayesian decoding of natural images from retinal neurons. In Advances in Neural Information Processing Systems, pages 6434–6445, 2017.

Adam Paszke, Sam Gross, Francisco Massa, Adam Lerer, James Bradbury, Gregory Chanan, Trevor Killeen, Zeming Lin, Natalia Gimelshein, Luca Antiga, et al. PyTorch: An imperative style, high-performance deep learning library. In Advances in Neural Information Processing Systems, pages 8024–8035, 2019.

Eftychios A Pnevmatikakis, Daniel Soudry, Yuanjun Gao, Timothy A Machado, Josh Merel, David Pfau, Thomas Reardon, Yu Mu, Clay Lacefield, Weijian Yang, et al. Simultaneous denoising, deconvolution, and demixing of calcium imaging data. Neuron, 89(2):285–299, 2016.

Pavlo M Radiuk. Impact of training set batch size on the performance of convolutional neural networks for diverse datasets. Information Technology and Management Science, 20(1):20–24, 2017.

Edward Raff. A Step Toward Quantifying Independently Reproducible Machine Learning Research. In Advances in Neural Information Processing Systems. Curran Associates, Inc., 2019. URL https://arxiv.org/pdf/1909.06674.pdf.

Matthew Rocklin. Dask: Parallel Computation with Blocked algorithms and Task Scheduling. In Proceedings of the 14th Python in Science Conference, pages 130 – 136, 2015.

Sergiu Sanielevici, Subhashini Sivagnanam, Kenneth Yoshimoto, Nicholas T Carnevale, and Amit Majumdar. The Neuroscience Gateway: Enabling Large Scale Modeling and Data Processing in Neuroscience. In PEARC ‘18: Proceedings of the Practice and Experience on Advanced Research Computing, page 52, 2018. ISBN 9781450364461. doi: 10.1145/3219104.3219139. URL `nsg`.

Shreya Saxena, Ian Kinsella, Simon Musall, Sharon H Kim, Jozsef Meszaros, David N Thibodeaux, Carla Kim, John Cunningham, Elizabeth MC Hillman, Anne Churchland, et al. Localized semi-nonnegative matrix factorization (LocaNMF) of widefield calcium imaging data. PLOS Computational Biology, 16(4):e1007791, 2020.

Caroline A Schneider, Wayne S Rasband, and Kevin W Eliceiri. NIH Image to ImageJ: 25 years of image analysis. Nature Methods, 9(7):671–675, 2012. ISSN 1548-7091. doi: 10.1038/nmeth.2089.

Jens F Schweihoff, Matvey Loshakov, Irina Pavlova, Laura Kuck, Laura A Ewell, and Martin K Schwarz. DeepLab-Stream: Closing the loop using deep learning-based markerless, real-time posture detec-tion. bioRxiv, 2019.

Christoph Sommer, Christoph Straehle, Ullrich Koethe, and Fred A Hamprecht. Ilastik: Interactive learning and segmentation toolkit. In 2011 IEEE international symposium on biomedical imaging: From nano to macro, pages 230–233, 2011.

Victoria Stodden, Jennifer Seiler, and Zhaokun Ma. An empirical analysis of journal policy effectiveness for computational reproducibility. Proceedings of the National Academy of Sciences, 115(11):2584–2589, 2018.

David Sussillo, Rafal Jozefowicz, L F Abbott, and Chethan Pandarinath. LFADS - Latent Factor Analysis via Dynamical Systems. arXiv, 2016.

Jeffery L. Teeters, Keith Godfrey, Rob Young, Chinh Dang, Claudia Friedsam, Barry Wark, Hiroki Asari, Simon Peron, Nuo Li, Adrien Peyrache, Gennady Denisov, Joshua H. Siegle, Shawn R. Olsen, Christopher Martin, Miyoung Chun, Shreejoy Tripathy, Timothy J. Blanche, Kenneth Harris, György Buzsáki, Christof Koch, Markus Meister, Karel Svoboda, and Friedrich T. Sommer. Neurodata Without Borders: Creating a Common Data Format for Neurophysiology. Neuron, 88(4):629–634, 2015. ISSN 0896-6273. doi: 10.1016/j.neuron.2015.10.025.

Michael Terry, Matthew Kay, and Ben Lafreniere. Perceptions and practices of usability in the free/open source software (FoSS) community. In Proceedings of the SIGCHI Conference on Human Factors in Computing Systems,pages 999–1008, 2010.

John Towns, Gregory D. Peterson, Ralph Roskies, J. Ray Scott, Nancy Wilkins-Diehr, Timothy Cockerill, Maytal Dahan, Ian Foster, Kelly Gaither, Andrew Grimshaw, Victor Hazlewood, Scott Lathrop, and Dave Lifka. XSEDE: Accelerating Scientific Discovery. Computing in Science & Engineering, 16(5):62–74, 2014. ISSN 1521-9615. doi: 10.1109/mcse.2014.80.

John W Tukey. Tukey1961.pdf. The annals of mathematical statistics, 33(1):1–67, 1962.

Joshua T Vogelstein, Brett Mensh, Michael Hausser, Nelson Spruston, Alan C Evans, Konrad Kording, Katrin Amunts, Christoph Ebell, Jeff Muller, Martin Telefont, et al. To the cloud! A grassroots proposal to accelerate brain science discovery. Neuron, 92(3):622–627, 2016.

David Waltz and Bruce G Buchanan. Automating science. Science, 324(5923):43–44, 2009.

Matthew R Whiteway, Dan Biderman, Yoni Friedman, Mario Dipoppa, E Kelly Buchanan, Anqi Wu, John Zhou, Jean-Paul Noel, The International Brain Laboratory, John Cunningham, and Liam Paninski. Partitioning variability in animal behavioral videos using semi-supervised variational autoencoders. bioRxiv, page 2021.02.22.432309, 2021. doi: 10.1101/2021.02.22.432309.

Alexander B Wiltschko, Matthew J Johnson, Giuliano Iurilli, Ralph E Peterson, Jesse M Katon, Stan L Pashkovski, Victoria E Abraira, Ryan P Adams, and Sandeep Robert Datta. Mapping sub-second structure in mouse behavior. Neuron, 88(6):1121–1135, 2015.

Anqi Wu, E Kelly Buchanan, Matthew R Whiteway, Michael Schartner, Guido Meijer, Jean-Paul Noel, Erica Rodriguez, Claire Everett, Amy Norovich, Evan Schaffer, Neeli Mishra, C Daniel Salzman, Dora Angelaki, Andráes Bendesky, The International Brain Laboratory, John Cunningham, and Liam Paninski. Deep Graph Pose: a semi-supervised deep graphical model for improved animal pose tracking. bioRxiv, page 2020.08.20.259705, 2020. doi: 10.1101/2020.08.20.259705.

Dimitri Yatsenko, Jacob Reimer, Alexander S. Ecker, Edgar Y. Walker, Fabian Sinz, Philipp Berens, Andreas Hoenselaar, R. James Cotton, Athanassios S. Siapas, and Andreas S. Tolias. DataJoint: managing big scientific data using MATLAB or Python. bioRxiv, 2015.

Andy B. Yoo, Morris A. Jette, and Mark Grondona. Job Scheduling Strategies for Parallel Processing, 9th International Workshop, JSSPP 2003, Seattle, WA, USA, June 24, 2003. Revised Paper. Lecture Notes in Computer Science, pages 44–60, 2003. ISSN 0302-9743. doi: 10.1007/10968987\_3.

Byron M Yu, John P Cunningham, Gopal Santhanam, Stephen I Ryu, Krishna V Shenoy, and Maneesh Sa-hani. Gaussian-process factor analysis for low-dimensional single-trial analysis of neural population activity. J Neurophysiol, 102(1):614–35, 2009. doi: 10.1152/jn.90941.2008.

Luyin Zhao and Fadi P Deek. Improving open source software usability. AMCIS 2005 Proceedings, page 430, 2005.

AC Zhou, B He, S Ibrahim, R Buyya, RN Calheiros, and AV Dastjerdi. eScience and Big Data Workflow in Clouds: A Taxonomy and Survey. Big data: Principles and paradigms, pages 431–456, 2016.

Pengcheng Zhou, Shanna L Resendez, Jose Rodriguez-Romaguera, Jessica C Jimenez, Shay Q Neufeld, Andrea Giovannucci, Johannes Friedrich, Eftychios A Pnevmatikakis, Garret D Stuber, Rene Hen, et al. Efficient and accurate extraction of in vivo calcium signals from microendoscopic video data. Elife, 7:e28728, 2018.

